# Flexible reprogramming of *Pristionchus pacificus* motivation for attacking *Caenorhabditis elegans* in predator-prey competition

**DOI:** 10.1101/2021.03.09.434602

**Authors:** Kathleen T. Quach, Sreekanth H. Chalasani

## Abstract

Animals with diverse diets must adapt their food priorities to a wide variety of environmental conditions. This diet optimization problem is especially complex for predators that compete with prey for food. Although predator-prey competition is widespread and ecologically critical, it remains difficult to disentangle predatory and competitive motivations for attacking competing prey. Here, we dissect the foraging decisions of the omnivorous nematode *Pristionchus pacificus* to reveal that its seemingly failed predatory attempts against *Caenorhabditis elegans* are actually motivated acts of efficacious territorial aggression. While *P. pacificus* easily kills and eats larval *C. elegans* with a single bite, adult *C. elegans* typically survives and escapes from bites. However, nonfatal biting can provide competitive benefits by reducing access of adult *C. elegans* and its progeny to bacterial food that *P. pacificus* also eats. We show that *P. pacificus* considers costs and benefits of both predatory and territorial outcomes to decide which food goal, prey or bacteria, should guide its motivation for biting. These predatory and territorial motivations impose different sets of rules for adjusting willingness to bite in response to changes in bacterial abundance. In addition to biting, predatory and territorial motivations also influence which search tactic *P. pacificus* uses to increase encounters with *C. elegans*. When treated with an octopamine receptor antagonist, *P. pacificus* switches from territorial to predatory motivation for both biting and search. Overall, we demonstrate that *P. pacificus* assesses alternate outcomes of attacking *C. elegans* and flexibly reprograms its foraging strategy to prioritize either prey or bacterial food.

## Introduction

Animals that exploit diverse food resources are more resilient to suboptimal environmental conditions than animals with specialized diets^1,2^. To fully benefit from versatile diets, animals must judge which food types and quantities maximize the ratio of energy intake to energy costs. Emphasis on calorie-rich and abundant foods is a sufficient strategy when foods are static and encountered one at a time^3^, but diet decisions are often conducted in more complex environments. For example, travel time to and between foods should also be minimized when multiple foods are simultaneously encountered^4,5^. When hunting mobile prey, predators should select prey that are easy to capture and pursue^6^. In addition to directly securing and eating foods, animals can indirectly prioritize foods by interfering with the ability of competitors to access those foods. However, little is known about the strategies that guide foragers when all these factors combine to produce a complex but naturalistic foraging problem in which a predator competes with prey for another food.

For predators that consume foods from different trophic levels, prey may consume and directly reduce the abundance of another of the predator’s food choices. This predator-prey competition (intraguild predation) is a widespread trophic motif in many food webs^7^, and its emergent effects on population dynamics and biodiversity remain widely researched and debated^8,9^. Here, killing prey simultaneously achieves dual food benefits by enabling consumption of prey corpses and reducing competition for shared food resources^8,10^. However, it is often unclear which of these predatory and competitive benefits is the dominant motivation for attacking competing prey. Evidence against predatory motivation comes from studies showing that corpses of competing prey are left uneaten more often than those of “true” prey that don’t compete with the predator^11,12,13,14^. Using similar logic to argue the opposite, studies that dismiss competitive motivation show that aggressive threat displays were absent against competing prey but frequently presented to intraspecific (non-prey) competitors^15^. However, killing prey without feeding can still indirectly promote predation^16^, and threat displays are not always needed in competitive fights^17^. To resolve these conflicting results and disentangle the motivations that drive a predator to attack a competing prey, more definitive and positive indicators are needed.

The predatory nematode *Pristionchus pacificus* (Figure 1A), its competing prey *Caenorhabditis elegans* (Figures 1B and 1C), and a shared bacterial food comprise a convenient laboratory system for investigating factors that influence an omnivorous predator’s diet decisions^18^. *P. pacificus* prefers to eat bacteria but can also use its teeth (Figures S1A and S1B) to attack and eat *C. elegans*^19,20^. In contrast, *C. elegans* lacks teeth (Figure S1C) and almost exclusively feeds on bacteria. To feed on *C. elegans, P. pacificus* must pierce the cuticle to access and ingest the internal pseudocoelomic fluid, thereby causing *C. elegans* death. While larvae are typically killed by a single bite (Figures 1D and 1E; Video S1), adult *C. elegans* are rarely killed and easily escape *P. pacificus* bites (Figures 1F and 1G; Video S2). Since nonfatal bites may be failed predatory attempts, this precludes the use of prey-feeding (which is only possible in successful predation) as a behavioral indicator of predatory motivation. Furthermore, nonfatal bites are executed similarly to bites that consummate in feeding, with no discernable threat displays that may suggest competitive motivation. Here, we deconstruct *P. pacificus* foraging decisions to show that nonfatal adult-targeted bites are not failed predatory attempts, but are instead goal-directed acts of aggression to expel competitors from a bacterial territory. Overall, we demonstrate that *P. pacificus* conducts cost-benefit analyses to flexibly switch between and adjust predatory and territorial strategies for biting *C. elegans*.

**Figure 1.**
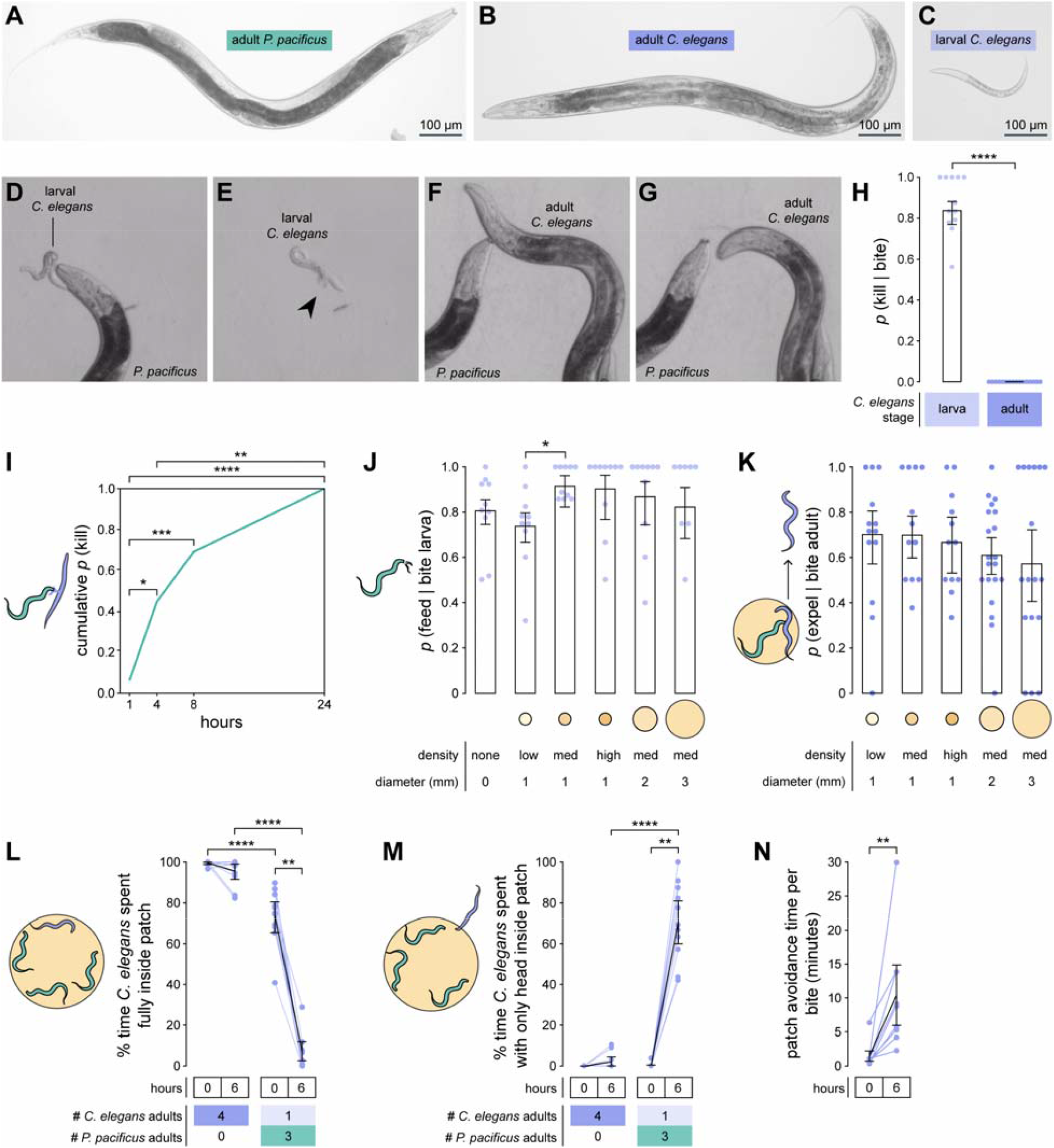
Nonfatal biting compels adult *C. elegans* to avoid bacteria occupied by *P. pacificus*. (A-C) Representative images of (A) young adult *P. pacificus*, (B) young adult *C. elegans*, and (C) larval *C. elegans* in L1 stage (earliest larval stage). (D-G) Images showing (D) *P. pacificus* biting larval (L1) *C. elegans*, (E) leakage of pseudocoelomic fluid after fatal biting of larval *C. elegans*, (F) *P. pacificus* biting adult *C. elegans*, and (G) adult *C. elegans* escaping from a bite. (H) Probability of *P. pacificus* killing different stages of *C. elegans* given a single bite (Wald test, n_*P*.*pacificus*_ = 12 – 16, n_bites per *P*.*pacificus*_ = 3 – 39). Individual data points are calculated as # kills / #bites for each *P. pacificus*. (I) Cumulative percentage of *P. pacificus* animals that successfully killed adult *C. elegans* by various time points (Fisher’s exact test with Benjamini-Hochberg adjustment, n_*P*.*pacificus*_ = 16). (J) Probability of feeding on bitten larva, across bacterial patch conditions (Wald test with single-step adjustment for Tukey contrasts, n_*p*.*pacificus*_ = 9-10, n_bites per *P. pacificus*_ = 1-31). (K) Probability that adult *C. elegans* exits a bacterial patch after being bitten, across bacterial patch conditions (Wald test with single-step adjustment for Tukey contrasts, n_*p*.*pacificus*_ = 13-20, n_bites per *P. pacificus*_ = 1-15). (L-M) Percentage of time that an adult *C. elegans* spent with its (L) body fully inside and (M) only its head inside a bacterial patch that contained either only *C. elegans* or three *P. pacificus* animals (Wilcoxon’s signed rank test (paired) and Dunn’s test (unpaired) with Benjamini-Hochberg adjustment, n_*C. elegans*_ = 11). (N) Average time that adult *C. elegans* spent avoiding a bacterial patch immediately after a bite (Wilcoxon’s signed rank test, n_*C. elegans*_ = 9). (H,J,K) Means and error bars are predicted probabilities and 95% confidence intervals from binomial logistic regression models of data. All other error bars are 95% bootstrap confidence intervals. * p<0.5, ** p<0.01, *** p<0.001, **** p<0.0001.

## Results

### Nonfatal biting compels adult *C. elegans* to avoid bacteria occupied by *P. pacificus*

To identify the different functions that *P. pacificus* may associate with biting, we first probed the immediate outcomes of biting *C. elegans* in the absence and presence of bacterial food. We assessed the ability of *P. pacificus* to kill *C. elegans* by confining them together in a small arena without any other food source, and then measuring how often individual bites resulted in fatality. We found that bites targeted at larval *C. elegans* mostly resulted in kills, while bites targeted at adult *C. elegans* rarely killed (Figures 1H and S1D). Even when allowed to focus all of its bites onto a single target, a single *P. pacificus* took ∼ 6 hours (Figures 1I and S1E) and ∼25 bites (Figure S1F) to kill adult *C. elegans*. Next, we analysed bites that occurred on bacterial patches (Figures S1G to S1L) to probe the potential use of biting for defending food territory. Most larva-targeted bites led to feeding on prey, regardless of whether bacteria were absent or present (Figures 1J and S1M). Since, *P. pacificus* biting rarely kills adult *C. elegans* and therefore rarely leads to prey-feeding, we instead monitored how often a bite led to adult *C. elegans* exiting a bacterial patch. The majority of adult-targeted bites that occurred on bacteria expelled adult *C. elegans* from a bacterial patch (Figures 1K and S1N; Video S3). Since successful predation also eliminates competition, larva-targeted biting simultaneously achieves both predatory and territorial benefits with relative ease. In adult-targeted biting, predation is rare and labor-intensive, but expulsion of intruders from bacteria can be achieved without killing.

To explore the long-term effects of nonfatal biting on adult *C. elegans* patch-leaving behavior, we placed adult *C. elegans* with or without *P. pacificus* for 6 hours on a small bacterial patch. With *P. pacificus* absent, adult *C. elegans* animals spent almost all of their time with their bodies fully inside the bacterial patch (Figure 1L). Upon initial exposure to *P. pacificus*, adult *C. elegans* still spent most of its time fully inside the lawn, though less than when *P. pacificus* was absent (Figures 1L and S2A). After 6 hours of predator exposure, *C. elegans* almost completely avoided fully entering the bacterial patch (Figures 1L), opting instead to insert only its head inside the patch (Figures 1M, S2B, and S2C). Additionally, the average time that adult *C. elegans* spent avoiding the lawn after each bite increased fivefold (Figure 1N), suggesting that adult *C. elegans* was conditioned at 6 hours to associate the bacterial patch with danger. Therefore, long-term nonfatal biting of adult *C. elegans* induces persistent avoidance of bacteria that is energetically efficient for *P. pacificus* to maintain.

### Progeny of predator-exposed adult *C. elegans* experience reduced access to bacteria

In order for nonfatal biting to have meaningful territorial benefits for *P. pacificus*, the relative fitness of *P. pacificus* would have to be higher than that of *C. elegans*. We speculated that biting-induced patch avoidance would force adult *C. elegans* to lay eggs away from bacteria. To test this, we developed an egg distribution assay (Figures 2A, and S2D to S2G) to measure where eggs were laid relative to a small bacterial patch over a 7-hour period (before eggs hatched). When *C. elegans* or *P. pacificus* adults laid eggs separately from each other, most eggs were deposited inside the patch, indicating absence of within-species territoriality (Figures 2B and 2C). In mixed groups, however, *C. elegans* was more likely to lay eggs off the patch as number of *P. pacificus* adults increased (Figure 2D), shifting the spatial distribution of *C. elegans* eggs away from the patch (likelihood ratio test on linear mixed-effects model, *χ*^*2*^ = 42.594, df = 3, *p* = 3.001e-09; Figures 2B, 2C, and 2E). Meanwhile, *P. pacificus* egg distribution was unaffected by the mix of adult *P. pacificus* and adult *C. elegans* (likelihood ratio test on linear mixed-effect model, *χ*^*2*^ = 5.1518, df = 3, *p* = 0.161; Figures 2B and 2C). The number of eggs laid per adult *C. elegans* remained unchanged (Figure S2H), indicating that *P. pacificus* rarely eats *C. elegans* eggs. Thus, biting interferes with *C. elegans* preference for laying eggs within bacteria.

**Figure 2.**
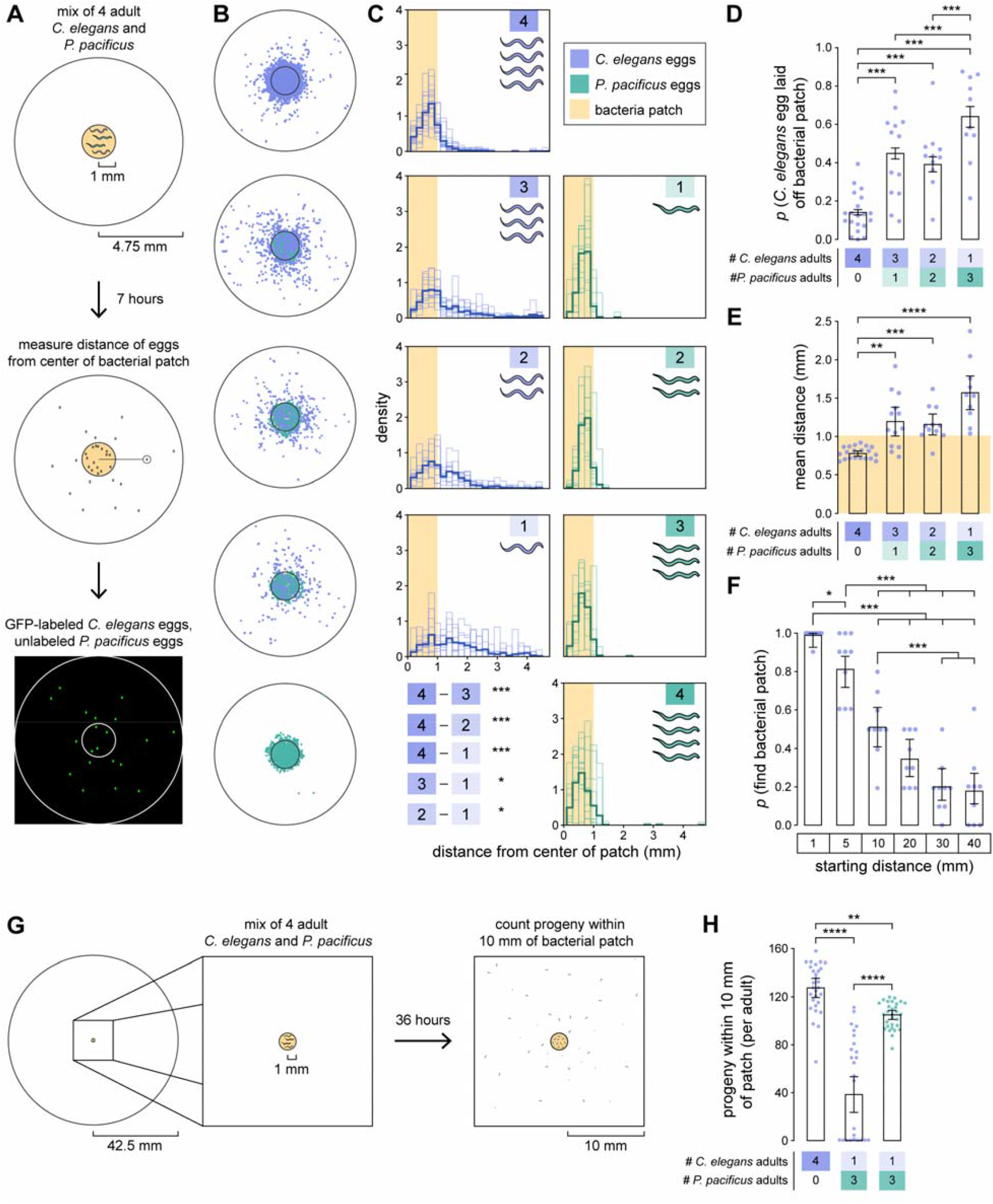
Progeny of predator-exposed adult *C. elegans* experience reduced access to bacteria. (A) Egg distribution assay. A mix of 4 adult *C. elegans* and *P. pacificus* laid eggs for 7 hours in an arena with a small bacterial patch. For each egg, species identity and distance from the center of the patch were determined. (B) Actual locations of eggs, pooled within the particular mix corresponding to same-row plots in (C). Outer concentric circle represents the arena bounds; inner concentric circle represents the bacterial patch. (C) Distributions of the radial distances of eggs laid by different mixes of adult *C. elegans* (left) and *P. pacificus* (right), relative to the center of a bacterial patch. Light-colored histograms represent egg distribution in individual arenas, while dark-colored histograms represent egg distribution pooled across all arenas. Light yellow shading indicates the radius (1 mm) of the bacterial patch. Egg distributions were compared within egg species (Wald test with single-step adjustment for Tukey contrasts, n_arena_ = 10 - 20). (D Percentage of *C. elegans* eggs that are laid off the bacterial patch (Wald test with single-step adjustment for Tukey contrasts, n_arena_ = 10 – 20, n_eggs per arena_ = 12 – 166). (E) Mean distance of *C. elegans* eggs for different mixes of adult *C. elegans* and *P. pacificus* (Dunn’s test, n_arena_ = 10-20). (F) Percentage of newly hatched larvae that find a small bacterial patch within 36 hours from various starting distances (Wald test with single-step adjustment for Tukey contrasts, n_arena_ = 9, n_larvae per arena_ = 10-11). (G) Egg distribution assay, with extended egg-laying time (36 hours) and increased maximum distance from the small bacterial patch (42.5 mm). Progeny within 10 mm of the patch were counted. (H) Number of *C. elegans* progeny (purple dots) and *P. pacificus* progeny (green dots), per adult, within 10 mm of a small patch (Dunn’s test, n_arena_ = 29-30). (D,F) Means and error bars are predicted probabilities and 95% confidence intervals from binomial logistic regression models of data. All other error bars are 95% bootstrap confidence intervals. * p<0.5, ** p<0.01, *** p<0.001, **** p<0.0001.

We next asked whether newly hatched *C. elegans* larvae would struggle to find a distant small bacterial patch. Since failure to find food within 36 hours induces arrested reproductive development in larval *C. elegans* (‘dauer’ state)^21^, we gently placed larvae (cleaned of bacteria) at various distances from a small bacterial patch and counted how many found the patch within 36 hours (see Methods: Patch-finding). At 10 mm starting distance away from the patch, larvae had only a ∼0.5 probability of finding the patch, with lower probabilities at greater starting distances (Figure 2F). To verify that biting causes adult *C. elegans* to lay eggs at these unfavorable distances, we spatially and temporally extended the egg distribution assay to 100 mm and 36 hours, respectively (Figure 2G). With *P. pacificus* present, the number of *C. elegans* larvae within 10 mm of the bacterial patch reduced to less than half of the number observed when predators were absent (Figure 2H). While *C. elegans* were typically more prolific than *P. pacificus* (Figure S2H), *P. pacificus* progeny outnumbered *C. elegans* progeny within 10 mm of the bacterial patch (Figure 2H). Altogether, these results illustrate how nonfatal biting accrues long-term territorial benefits and increases *P. pacificus* fitness relative to that of *C. elegans*.

### *P. pacificus* inflicts non-fatal biting to achieve territorial outcomes

While we have thus far shown that nonfatal biting of adult *C. elegans* provides territorial benefits, it remains to be shown whether these territorial benefits are goal-directed or a serendipitous consolation prize of botched predation. To assess food values, we first analysed at the long-term net energy yield of various single-food diets. *P. pacificus* were allowed to feed freely on excess bacteria (*E. coli* OP50), larval *C. elegans*, or adult *C. elegans* for 6 hours before being stained for fat stores with the lipophilic dye Oil Red O (see Methods: Oil Red O staining). *P. pacificus* fed with bacteria displayed the most stained fat, followed by adult-fed and then larva-fed *P. pacificus* (Figures 3A and S3A to S3D). Thus, bacteria-based diets are associated with higher energy yields than prey-based diets. Further, given enough time for successful predation, a diet comprised of adult *C. elegans* is more efficient than a larva-based diet.

**Figure 3.**
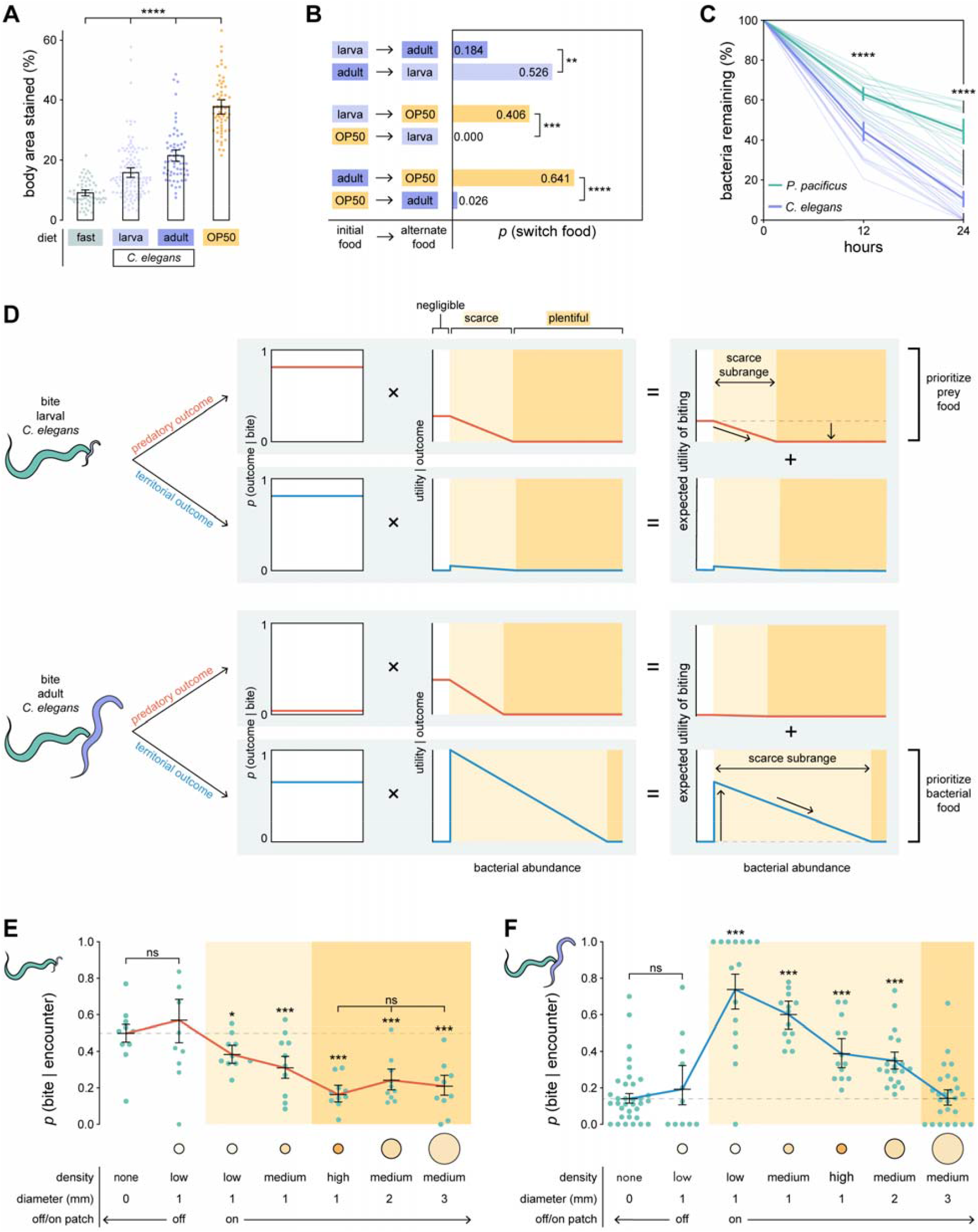
*P. pacificus* inflicts non-fatal biting to achieve territorial outcomes. (A) Percentage of *P. pacificus* body stained with Oil Red O after 6 hours on different diets (Dunn’s test with Benjamini-Hochberg adjustment, n_*P. pacificus*_ = 60-117). (B) Probability of *P. pacificus* switching from one food source to an alternate food source (Fisher’s exact test with Benjamini-Hochberg adjustment, n_*P. pacificus*_ = 29-39). (C) Percentage of a bacterial patch remaining after placing a single adult *C. elegans* or *P. pacificus* on the patch (Dunn’s test with Benjamini-Hochberg adjustment, n_adult_ = 16-23). (D) Model of how biting valuation should adjust to different *C. elegans* targets and bacterial abundances. Expected utility of biting is calculated as the sum of the utilities (subjective values) of potential biting outcomes (predatory or territorial), weighted by their respective probabilities. Therefore, expected utility of biting has both a predatory component and a territorial component. White, light yellow, and dark yellow shading represent negligible, scarce, and plentiful subranges, respectively, of bacterial abundance. Dashed lines represent expected utility of biting when bacteria abundance is negligible, with vertical arrows indicating predicted direction of change in expected utility of biting. Sloped arrows represent predicted monotonic decreases in expected utility of biting. Bacterial abundance and utility are in arbitrary units. (E) Probability that *P. pacificus* bites given an encounter with larval *C. elegans* (Wald test with single-step adjustment for Tukey contrasts, n_*P. pacificus*_ = 9-10, n_encounters per *P. pacificus*_ = 1-66. Red line highlights similarity to the shape predicted for predatory biting in (D). (F) Probability that *P. pacificus* bites given an encounter with adult *C. elegans* (Wald test with single-step adjustment for Tukey contrasts, n_*P. pacificus*_ = 12-34, n_encounters per *P. pacificus*_ =1-38). Blue line highlights similarity to the shape predicted for territorial biting in (D). (E,F) Means and error bars are predicted probabilities and 95% confidence intervals from binomial logistic regression models of data. All other error bars are 95% bootstrap confidence intervals. ns p>0.05, * p<0.5, ** p<0.01, *** p<0.001, **** p<0.0001.

Although adult prey is more valuable than larval prey in the long term, *P. pacificus* may discount delayed rewards^22^ to avoid food deprivation. To assess short-term food preference, we placed *P. pacificus* in one of two plentiful, neighboring food patches and then checked whether *P. pacificus* switched to the alternative food patch after an hour (see Methods: Food switching). By comparing switching probabilities, we found that *P. pacificus* prefers bacteria over all prey, and larval prey over adult prey (Figure 3B). Preference for bacteria over larvae is consistent with previous findings^20^. Contrary to long-term food value (Figure 3A), *P. pacificus* preferred larval over adult prey (Figure 3B). Notably, inverse switches (a→b, b→a) had combined probabilities less than 1 (Figure 3B), which is consistent with previous reports that nematodes tend to stay within a food patch^23^, and with foraging theory that discounts food value by the time it takes to travel to food^4,5^. Overall, preference for easily consumed foods and for closer foods suggest that *P. pacificus* prefers immediate over delayed food rewards.

We next explored the potential competitive benefits associated with the territorial outcome of biting. To compare their ability to exploit bacteria, we placed an adult *C. elegans* and an adult *P. pacificus* onto separate identical patches of GFP-labelled bacteria, and then measured bacterial fluorescence at 12 and 24 hours. We found that adult *C. elegans* consumed bacteria ∼1.5x faster than adult *P. pacificus* at both time points (Figure 3C). Eggs laid by adult *C. elegans* began hatching at 12 hours, with a range of 20 to 62 larvae present by 24 hours (Figures S3E and S3F). However, bacterial consumption rate does not increase between 12 and 24 hours (Figure 3C), and we found no correlation between number of larvae and bacteria consumed (Pearson’s *r* = 0.2480, *p* = 0.8066). These results show that adult *C. elegans* more efficiently exploits bacteria and may outcompete *P. pacificus* for bacteria, but larvae pose a negligible short-term competitive threat.

To determine the relative contributions of predatory and territorial outcomes toward the overall subjective value of biting a particular stage of *C. elegans* target (larva or adult), we applied neuroeconomic theories of how to make rational decisions about actions that have probabilistic outcomes. In expected utility theory^24^, the expected utility (overall subjective value) of an action takes into account both the probability that a particular outcome will occur given an action, in addition to the utility (subjective value) attained if that outcome occurs. The expected utility of an action for a particular context is calculated as the sum of the utilities (subjective values) of each outcome, weighted by their respective probabilities of occurring given an action. To calculate the expected utility of biting larval or adult *C. elegans*, we would have to first determine the probability that predatory and territorial outcomes occur given a bite, as well as how utility of those outcomes changes with bacterial abundance.

First, we contrived food choices such that *P. pacificus* only encounters larval *C. elegans* or only adult *C. elegans*, so that the decision is between biting outcomes, rather than between different prey options (Figure 3D). We then assigned probabilities to each biting outcome, for each type of *C. elegans* target (Figure 3D). After biting either larval or adult *C. elegans*, we assumed that a bite can lead to two possible outcomes: the predatory outcomes leads to feeding on prey, while the territorial outcome eliminates competitors for bacterial food (Figure 3D). For biting larval *C. elegans*, we set both *p(predatory outcome*|*bite)* and *p(territorial outcome*|*bite)* as equal to the pooled probability that a bite leads to feeding on larva (*p(feed*|*bite)*= 0.8115; Figure S1M), since feeding on larvae simultaneously eliminates competitors (Figure 3D). For biting adult *C. elegans*, we assigned *p(predatory outcome*|*bite)* a very low probability (see Methods: Expected utility of biting) since adult *C. elegans* is rarely killed by a *P. pacificus* bite (Figures 1H, 1I, and S1F). In contrast, we assigned *p(territorial outcome*|*bite)* as equal to the pooled probability that a bite expels adult *C. elegans* from a bacterial patch (*p(expel*|*bite) =* 0.6483; Figure S1N).

Next, we determined how the utility of predatory or territorial outcomes changes with bacterial abundance. We subdivided bacterial abundance into three behaviorally defined subranges: 1) in the ‘negligible’ subrange, *P. pacificus* considers bacteria too meager or absent to exploit, 2) in the ‘scarce’ subrange, *P. pacificus* exploits bacteria, but must bite to increase food supply, and 3) in the ‘plentiful’ subrange, *P. pacificus* has excess bacterial food and therefore does not need to bite to secure supplementary food. The bounds of these subranges should shift depending on whether the intended biting outcome is predatory or territorial, and on whether the *C. elegans* target is larva or adult. In particular, we are interested in how outcome values in the scarce subrange compare to those in the negligible and plentiful subranges.

We first described the general shape of how the utility of predatory biting outcomes (for both larval and adult *C. elegans* targets) should change with bacterial abundance (Figure 3D). Since the goal of predation is to kill prey for food, the value of predatory biting outcomes should be highest when bacterial abundance is negligible, and then monotonically decrease as the abundance of its preferred food, bacteria, increases (Figure 3D). This is consistent with previous reports that *P. pacificus* bites larvae less when bacteria are present than when absent^20^. We used ORO fat-staining of prey relative to bacteria (Figure 3A) to estimate predatory biting value over the negligible subrange, and probabilities of switching from prey to bacteria (Figure 3B) to estimate the rate at which predatory value degrades over the scarce subrange as bacterial abundance increases (see Methods: Expected utility of biting). By definition, the plentiful subrange begins where the utility of predatory biting outcomes reaches zero. Even though the utility of predatory biting outcomes is higher for adult prey than for larval prey over the negligible subrange, *P. pacificus* should drop adult prey from its diet at a lower bacterial abundance than for larval prey due to preference for immediate food rewards (Figure 3D).

In contrast to the utility of predatory biting outcomes, we characterized the utility of territorial biting outcomes (for both larval and adult *C. elegans* targets) as generally having a non-monotonic shape and a sharp peak (Figure 3D). Under the goal of territoriality, biting should remove competitors from a bacterial territory to indirectly prioritize bacterial food. Over the negligible subrange, bacteria are absent or not worth defending, and therefore the utility of territorial outcomes should be zero (Figure 3D). As the negligible subrange transitions to the lower bound of the scarce subrange, scarcity-induced competitive pressure is strongest and should induce a sudden peak in the utility of territorial outcomes (Figure 3D). From there, utility of territorial outcomes should monotonically decrease as bacterial abundance increases (Figure 3D). This territorial value function is similar to previous energy cost models of feeding-based territoriality^25^. We used *C. elegans* consumption rate of bacteria relative to that of *P. pacificus* (Figure 3C) to estimate peak territorial biting values (see Methods: Expected utility of biting). Additionally, we predicted that the territorial scarce subranges would be wider than predatory scarce subranges, due to bacterial loss to competitors (Figure 3D, see Methods: Expected utility of biting). Accordingly, the territorial plentiful subrange begins at a higher bacterial abundance compared to predatory plentiful subrange, and represents excess bacterial abundance that accommodates both *P. pacificus* and adult *C. elegans* (Figure 3D). Notably, predatory and territorial value functions have similar shapes across scarce and plentiful subranges, but differ in whether the value over the negligible subrange is higher or lower than adjacent values in the scarce subrange (Figure 3D). It is important to note that the model in Figure 3D only specifies the shape of biting incentive by predicting 1) the direction of change across bacterial abundance subranges, particularly compared to when bacteria is negligible, 2) monotonic decrease across the scarce subrange, 3) higher peak expected utility of biting for adult *C. elegans* target, and 4) wider scarce subrange for biting of adult *C. elegans*.

For each combination of outcome type (predatory or territorial) and *C. elegans* target (larval or adult), we multiplied the outcome probability, *p(outcome*|*bite)*, by its corresponding outcome utility function (*utility*|*bite*) (Figure 3D). The expected utility of biting a particular stage of *C. elegans* is then estimated as the sum of the probability-weighted utilities for both predatory and territorial outcomes, such that expected utility of biting as both a predatory component and a territorial component (Figure 3D). Therefore, the outcome with the higher probability-weighted utility should be the primary contributor towards biting motivation. For biting larval *C. elegans*, the predatory outcome should be prioritized due to low competitive pressure from larvae (Figure 3D). For biting adult *C. elegans*, the territorial outcome should be prioritized due to the low probability of killing adult prey (Figure 3D).

Next, we tested our predictions to determine whether *P. pacificus* considers both predatory and territorial outcomes, or only predatory outcomes, to make goal-directed biting choices. We used the probability that *P. pacificus* bites *C. elegans* given an encounter, *p(bite*|*encounter)*, to quantify biting incentive. We placed *P. pacificus* in an arena with either larval or adult *C. elegans*, and varied bacterial abundance by changing the size or density of a bacterial patch (Figures S1G to S1L). We then used the shape of how biting incentive changed with bacterial abundance to infer whether prey or bacteria are the intended food goal of biting. As predicted, we observed that larva-targeted biting incentive monotonically decreased as bacterial abundance increased (Spearman’s ρ = -0.58, *p* < 0.001; Figure 3E), resembling the predicted shape of the predatory component of expected utility of biting larval *C. elegans* (Figure 3D). By contrast, adult-targeted biting incentive was low when bacteria were negligible and high when bacteria were scarce (Figure 3F), which conforms with the predicted non-monotonic shape (Spearman’s ρ = 0.08, *p* = 0.374; Hoeffding’s *D* = 0.03, *p* < 0.001) of the territorial component of the expected utility of biting adult *C. elegans* (Figure 3D). However, adult-targeted biting exhibited monotonic decrease with increasing bacterial abundance when only on-patch bacterial conditions were considered (Spearman’s ρ = -0.74, *p* < 0.001; Figure 3F). Importantly, adult-targeted biting incentive diminishes at a higher bacterial abundance than larva-targeted biting incentive (Figure 3E and 3F), consistent with our prediction of a wider scarce subrange for adult-targeted territorial value. To examine the critical bacterial abundance threshold between negligible and scarce subranges (Figure 3D), we used a low-density bacterial patch (Figure S1H) that induced patch-staying and patch-leaving with roughly equal probabilities (Figure S4A, and Videos S4 and S5). Using a choice variability approach for probing decision-making^26^, we segregated encounters into off- and on-patch events to reflect *P. pacificus* decision to ignore or exploit the patch, respectively. Off- and on-patch biting incentive were not significantly different for larva-targeted bites (Wald test with single-step adjustment for Tukey contrasts, *p* = 0.07663; Figure 3E), while on-patch biting incentive was higher for adult-targeted bites (Figure 3F). Collectively, these results show that *P. pacificus* considers both predatory and territorial biting outcomes in a context-specific manner, and incorporates both *C. elegans* and bacterial information to direct predatory attacks against larval *C. elegans* and territorial aggression against adults.

### Territorial biting is driven by chemosensation and mechanosensation of bacteria

We next explored how *P. pacificus* senses bacteria for adjusting territorial biting incentive. While predatory biting could be suppressed only by increasing the density of smallest-sized bacterial patch (1 mm) (ANOVA; density: *F* = 17.84, df = 3, p < 0.0001; diameter: *F* = 22.19, df = 2, p = 0.168; Figure 3E), suppression of territorial biting required increasing both density and diameter of the bacterial patch (ANOVA; density: *F* = 11.668, df = 3, p < 0.0001; diameter: *F* = 1.838, df = 2, p < 0.0001; Figure 3F). This suggests that *P. pacificus* senses at least two features of bacteria that are relevant for territorial biting decisions. To identify a role for chemosensation, we ablated bilateral pairs of amphid neurons that have exposed cilia at the *P. pacificus* nose (Figure 4A). Relative to mock controls, ablation of ASH and AWC neurons decreased biting incentive against adult *C. elegans* on a medium-density, 1 mm bacterial patch, while ablation of ADL neurons increased biting incentive (Figure 4B). Although these neurons are poorly understood in *P. pacificus*, studies of homologous neurons in *C. elegans* do offer some clues. The olfactory neuron AWC triggers local search behavior upon removal from a bacterial patch^27^, and also senses bacteria-related odorants^28,29^. In *C. elegans*, ASH and ADL are involved in avoiding high ambient oxygen^30,31^ and migrating to the thick boundary of a bacterial patch, where local oxygen concentration is lower due to higher bacterial metabolism^27^. Recent studies report that *P. pacificus* can similarly distinguish between oxygen levels^32^ (albeit via different molecular mechanisms^33^), and mutants with cilia-defective amphid neurons exhibit impaired oxygen responses^34^.

**Figure 4.**
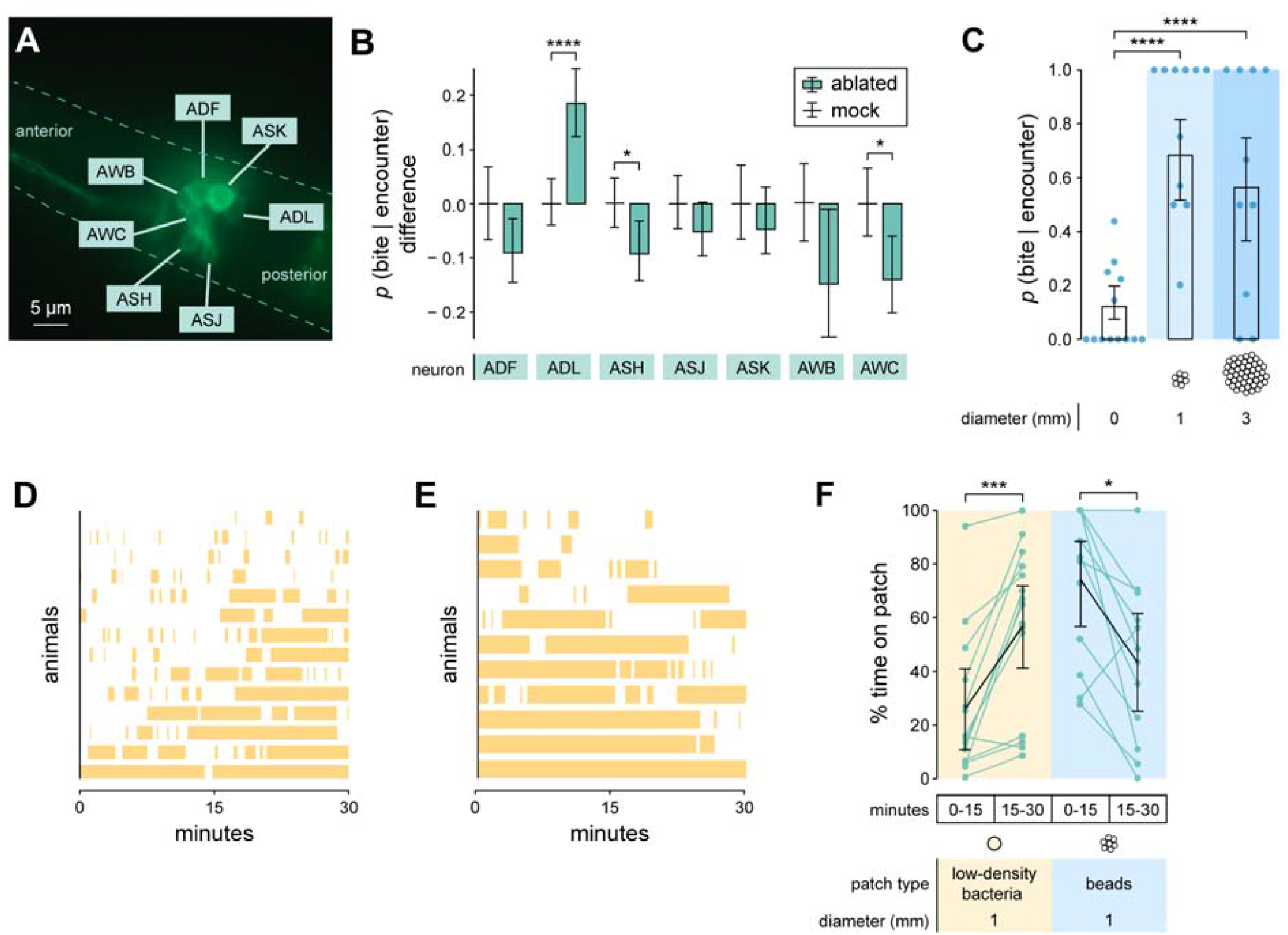
Territorial biting is driven by chemosensation and mechanosensation of bacteria. (A) DiO-stained chemosensory amphid neurons of *P. pacificus*. Dashed line demarcates head silhouette. (B) Difference in *p(bite*|*encounter)* between mock-ablated (mean centered at zero) and neuron-ablated *P. pacificus*, with an adult *C. elegans* target on a scarce bacterial lawn (Wald test with Benjamini-Hochberg adjustment, n_*P. pacificus*_ = 5-31, n_encounters per *P. pacificus*_ =2-27. (C) Probability that *P. pacificus* bites given an encounter with adult *C. elegans* on a patch of Sephadex beads (Wald test with single-step adjustment for Tukey contrasts, n_*P. pacificus*_ = 7-13, n_encounters per *P. pacificus*_ = 1-14. (D, E) Timecourse of *P. pacificus* residence on a 1 mm patch consisting of (D) low-density bacteria or (E) Sephadex beads. (F) Change in *P. pacificus* patch residence time for edible (low-density bacteria) and inedible (beads) low-residence patches (Wilcoxon’s signed rank test with Benjamini-Hochberg adjustment, n_*P. pacificus*_ = 11-14). (B,C) Means and error bars are predicted probabilities and 95% confidence intervals from binomial logistic regression models of data. All other error bars are 95% bootstrap confidence intervals. * p<0.5, ** p<0.01, *** p<0.001, **** p<0.0001.

In addition to chemosensation, we also probed how mechanosensation of bacteria modulates territorial biting. We measured adult-targeted biting incentive on patches composed of Sephadex gel beads, whose surfaces elicit mechanosensation similar to that of bacterial surfaces^35^. Importantly, these beads are inedible and lack the chemical signatures of live bacteria. As with low-density bacterial patches, *P. pacificus* spent less time on a bead patch than on medium- or high-density bacterial patches (Figure S4A). On-patch biting incentive was high (Figure 4C), similar to that associated with a low-density bacterial patch of the same size (Figure 3F). This high biting incentive was not suppressed in larger bead patches (Figure 4C), suggesting that increased patch size is insufficient to suppress biting incentive without some other bacterial sensory cue. *P. pacificus* decreased its residence time on a bead patch over time, opposite of what it does on a low-density bacterial patch (Figures 4D to 4F, and S4B), suggesting that eating is required to sustain patch exploitation. These results suggest that *P. pacificus* senses some minimal ‘bacterial’ abundance in these bead patches. Based on existing information about *P. pacificus* and its relative *C. elegans*, we surmise that sensation of bacterial odor may be used to locate a bacterial patch from afar, oxygen sensation may be used to locate the thick (oxygen-consuming) boundary of a bacterial patch, and mechanosensation may be used to detect low-density bacteria when odor and oxygen gradients are too low.

### Predatory and territorial biting are associated with different search tactics

To further confirm that predatory and territorial motivations for biting are distinct and separable, we tested how those motivations differentially guide how *P. pacificus* searches for *C. elegans* while simultaneously exploiting a scarce bacterial patch. We first contemplated how biting motivation should influence search speed. Under predatory motivation, *P. pacificus* should minimize search costs since prey are inferior to bacteria, and there is no urgency to hunt while prey reside on the patch. Rather than increase search speed, *P. pacificus* should graze normally on bacteria and opportunistically bite and feed on prey during chance encounters. Under territorial motivation, *P. pacificus* should swiftly find and expel intruders to halt rapid loss of preferred food to adult *C. elegans*. To test these predictions, we tracked *P. pacificus*’s location on a scarce bacterial patch with either *P. pacificus* or *C. elegans* cohabitants. Since both nematode species are attracted to the thick boundary of a bacterial patch^36,32^, we assessed *P. pacificus’s* incentive to search for *C. elegans* by measuring forward angular movement along the patch circumference (patrol speed, Video S6) relative to x-y movement (translational speed) (Figure 5A). We found that *P. pacificus* exhibited faster patrol and translational speed with adult *C. elegans* cohabitants than with larvae or with other *P. pacificus* (Figure 5B to 5E). Translational speed was slower with larvae than with *P. pacificus* (Figure 5E), likely due to stationary bouts of *P. pacificus* feeding on larvae. To discount these stationary bouts in speed considerations, we analyzed at the ratio of patrol speed to translational speed. This speed ratio was higher with adult *C. elegans*, signifying that *P. pacificus* directed a greater proportion of its locomotion to patrolling the patch boundary when adult *C. elegans* was present (Figure 5F). Speed ratio did not differ between patches with larvae and patches with only *P. pacificus* (Figure 5F), suggesting that grazing-style feeding on bacteria motivated both exploration patterns. Moreover, *P. pacificus* reduced its speed and speed ratio in contexts where *C. elegans* was conditioned to mostly avoid the bacterial patch and pose little competitive threat (Figure S5). While both larva-targeted and adult-targeted biting incentives were relatively high for the tested scarce bacterial patch, patrolling only increased with adult cohabitants, indicating that increased speed is not associated with biting in general. Thus, territorial but not predatory biting motivation increases patrolling speed, reflecting an active and energy-intensive search tactic that is commensurate with protecting an energy-rich bacterial food supply.

**Figure 5.**
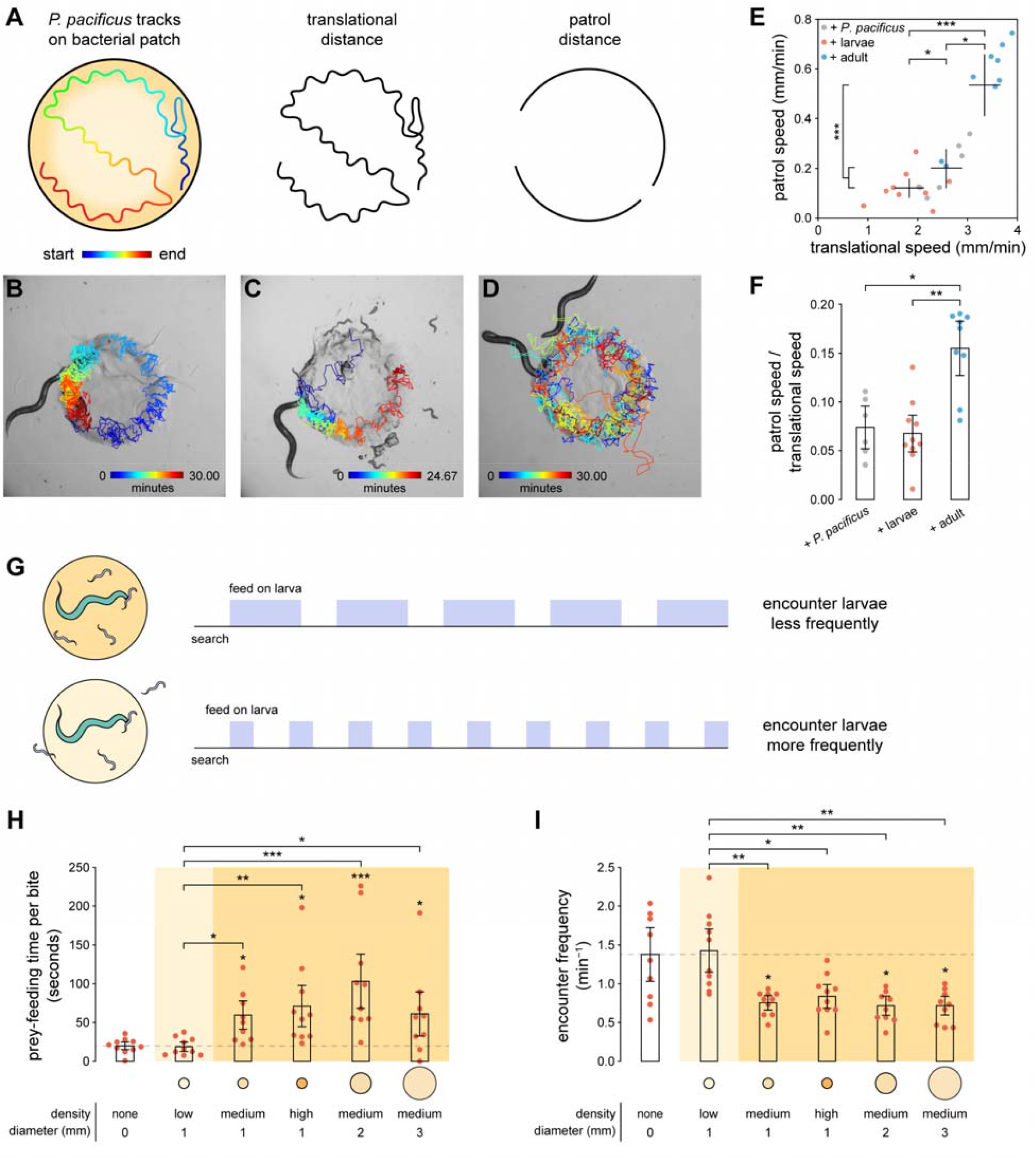
Predatory and territorial biting are associated with different search tactics. (A) Location of *P. pacificus* mouth was tracked during exploration of a bacterial patch. Translational distance was calculated as the total x-y distance traveled. Patrol distance was calculated as the forward radial distance traveled while *P. pacificus* explores the patch boundary. (B-D) Tracks of *P. pacificus* mouth while cohabiting a patch with (B) another *P. pacificus*, (C) larval *C. elegans*, and (D) adult *C. elegans*. (E) Translational and patrol speeds of *P. pacificus* with different cohabitants (Tukey’s HSD test, n_*P. pacificus*_ = 6-10). (F) Patrol speed as a proportion of translational speed, to discount paused exploration while feeding on larvae (Dunn’s test with Benjamini-Hochberg adjustment, n_*P. pacificus*_ = 6-10. (G) Prey-feeding duration was expected to be longer on bacteria-rich patches (top) compared to bacteria-poor patches (bottom). (H,I) Light and dark yellow shading indicates low- and high-residence patches, respectively (See fig. S7A). (H) Average prey-feeding time per bite (Dunn’s test with Benjamini-Hochberg adjustment, n_adult_ = 10). (I) Frequency of *P. pacificus* encounter with larval *C. elegans* (Dunn’s test with Benjamini-Hochberg adjustment, n_*P. pacificus*_ = 10). All error bars are 95% bootstrap confidence intervals. * p<0.5, ** p<0.01, *** p<0.001, **** p<0.0001.

We next asked if *P. pacificus* could engage alternate energy-efficient search tactics for increasing encounters with larval prey. *P. pacificus* must cease feeding on the current prey to resume search for other prey, so we reasoned that one way to increase prey encounter frequency without increasing search cost is to reduce prey-feeding time (Figure 5G). Prey are mobile, so while *P. pacificus* feeds on one prey, other prey could be dispersing, especially when bacterial abundance is not high enough to retain prey on the patch. To limit dispersal, *P. pacificus* can kill prey to immobilize it now and then finish eating later. To test this prediction, we measured the time *P. pacificus* spent feeding on larval prey (∼100 in each arena) across various types of bacterial patches. Importantly, larvae were able to escape the arena, making dispersal a real threat of food loss. As expected, we found that prey-feeding time per bite was lower when bacteria were absent or low-density, and higher when bacteria were medium- or high-density (Figure 5H). If prey-feeding time were closely associated with biting in general rather than with predatory strategy, then we would expect prey-feeding times that were graded with biting incentive. However, we did not see any difference between prey-feedings times on and off a low-density patch (*p* = 0.90558), or between a medium- and high-density 1 mm diameter patch (*p* = 0.99965) (Figure 5H), where we observed significant differences in biting incentive (Figure 3E). Instead, feeding times matched *P. pacificus*’s own patch-leaving behavior (Figure S1A), which may serve as a heuristic for judging other nematodes’ patch-leaving proclivity. Furthermore, we found that bacteria-free and low-density bacterial conditions associated with low prey-feeding times were also associated with more frequent encounters with prey, even though larvae were more dispersed (Figure 5I). Therefore, predatory motivation modulates prey-feeding time to implement a passive search tactic that is appropriate for the lower energy content of prey food. Overall, by using its biting motivation to coordinate search tactics, *P. pacificus* ensures that efforts are unified into a cohesive predatory or territorial foraging strategy.

### Blocking dopamine D2 or octopamine receptors modulates territoriality

We explored potential signaling mechanisms for regulating both the biting and search components of the territorial foraging strategy. Since knowledge of *P. pacificus* pathways is limited, we consulted known pathways in its well-researched relative, *C. elegans*. In *C. elegans*, the absence of bacteria attenuates D2-like receptor signaling (the biological action of amisulpride), which in turn triggers release of octopamine^37^, the invertebrate homolog of norepinephrine. We hypothesized that a similar pathway used for detecting bacterial scarcity may also exist in *P. pacificus* for modulating territorial behavior. Using a pharmacological approach, we exogenously treated *P. pacificus* with various compounds by dispensing a small volume of a concentrated drug solution onto a small bacterial patch (see Methods: Drug treatment), and then allowing *P. pacificus* to reside in that patch for two hours immediately before testing behavior. We found that treatment with dopamine D2 receptor antagonist, amisulpride, enhanced biting incentive when bacteria were scarce, but had no effect when bacteria were absent or plentiful (Figure 6A). This suggests that blocking D2 receptors does not affect general arousal to bite, but is context specific to conditions that signal competitive pressure. In contrast, treatment with epinastine, a high-affinity octopamine receptor antagonist^38^, affect biting incentive on all bacterial conditions tested (Figure 6B). Epinastine treatment affected biting incentive in opposite ways, depending on whether bacteria were absent or present. Specifically, epinastine treatment increased biting incentive when bacteria were absent and decreased it when bacteria were scarce or plentiful (Figure 6B). The overall result of epinastine treatment is that biting incentive monotonically decreased with bacterial abundance (Figure 6B), which is indicative of *P. pacificus* prioritizing predatory outcomes of biting (see Figure 3D). Moreover, epinastine treatment suppressed the increased patrolling associated with territorial biting (Figures 6C and 6D), suggesting that epinastine also affects territorial search tactics. While treatment with specific receptor antagonists affected territorial behavior, we did not see any change with treatment with a D2 receptor agonist or octopamine (Figure S6). Collectively, we found that blocking dopamine D2 receptors enhanced territorial biting, while blocking octopamine receptors induced *P. pacificus* to switch from a territorial to a predatory foraging strategy for biting adult *C. elegans*.

**Figure 6.**
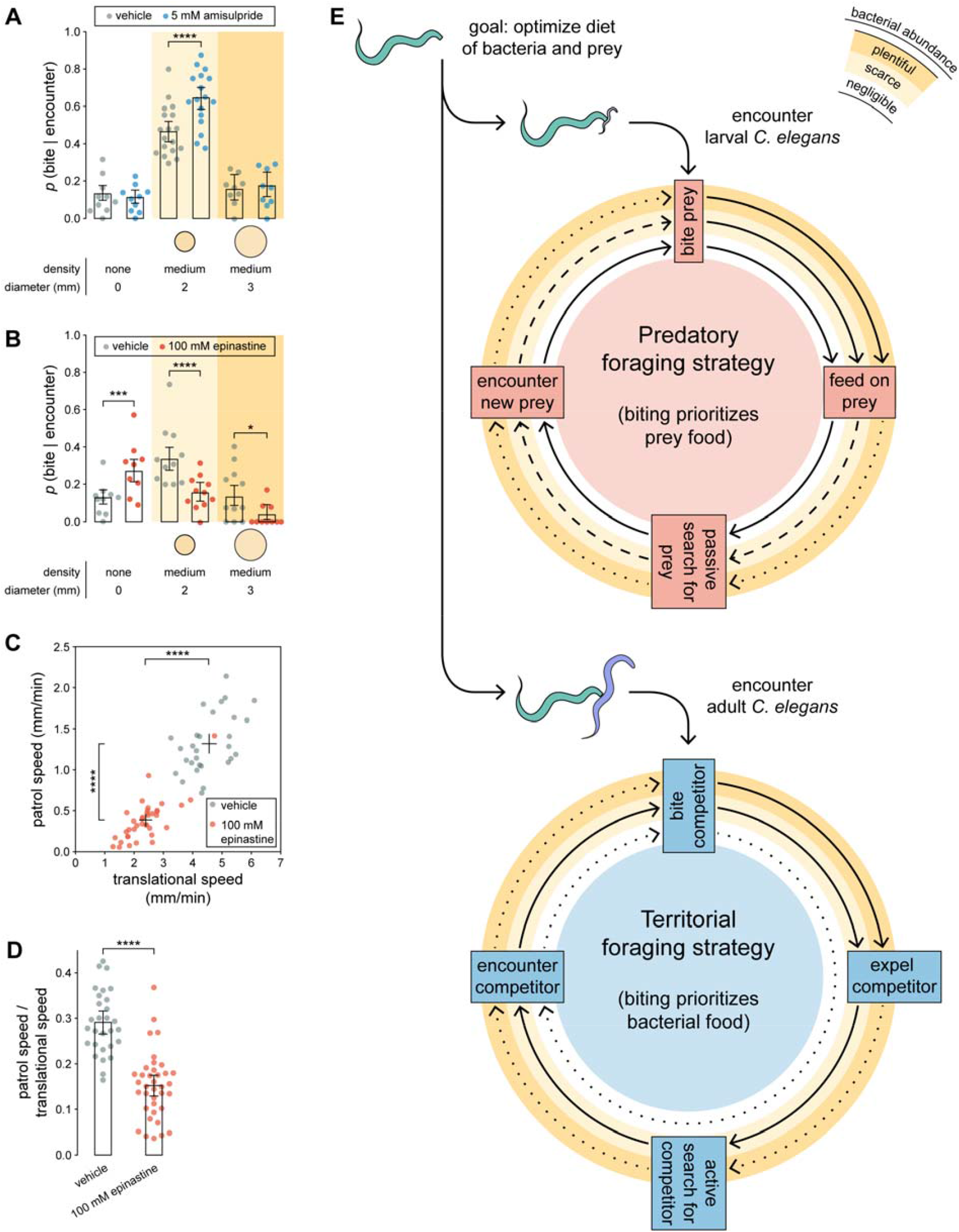
Blocking dopamine D2 or octopamine receptors modulates territorial biting. (A,B) *p(bite*|*encounter)* for *P. pacificus* treated with (A) the dopamine D2 receptor antagonist, amisulpride (Wald test with Benjamini-Hochberg adjustment, n_*P. pacificus*_ = 9-18, n_encounters per *P. pacificus*_ = 24-42), and (B) the octopamine receptor antagonist, epinastine (Wald test with Benjamini-Hochberg adjustment, n_*P. pacificus*_ = 9-11, n_encounters per *P. pacificus*_ = 9-38). (C,D) Effect of epinastine on (C) translational and patrol speeds, and on (D) patrol speed as a proportion of translational speed (Dunn’s test, n_*P. pacificus*_ = 29-37). (E) *P. pacificus* selects the predatory foraging strategy against larval *C. elegans*, and selects the territorial foraging strategy against adult *C. elegans*. Each foraging strategy consists of repeating cycles of four stages: bite → biting outcome → search for *C. elegans* → encounter *C. elegans*. Dark yellow, light yellow, and white concentric circles represent plentiful, scarce, and negligible bacterial abundance, respectively. Solid, dashed, and dotted arrows indicate highest, intermediate, and lowest probabilities, respectively, of achieving the next step. (A-D) Drug concentrations refer to the concentration of drug solution applied to a bacterial patch for treatment (see Methods: Drug treatment). (A,B) Means and error bars are predicted probabilities and 95% confidence intervals from binomial logistic regression models of data. All other error bars are 95% bootstrap confidence intervals. * p<0.5, ** p<0.01, *** p<0.001, **** p<0.0001.

## Discussion

We present a model of two distinct, flexible, and coordinated foraging strategies that *P. pacificus* uses for biting its competing prey, *C. elegans* (Figures 6E). *P. pacificus* engages the predatory foraging strategy (Figure 6E) against larval *C. elegans*, which is easy to kill and poses minimal competitive threat to the bacterial supply. Here, the goal of biting is to kill and eat prey, so *P. pacificus* bites most when prey is the only food source, and bites less as bacteria becomes more abundant. While most bites consummate in feeding on larval prey, *P. pacificus* can cut prey-feeding short and instead use that time to passively search for and immobilize larvae before they disperse. In contrast to the predatory strategy, the territorial strategy (Figure 6E) is activated against adult *C. elegans*, which is difficult to kill and consumes bacteria faster than *P. pacificus*. Instead of biting to acquire prey, here biting is used to protect valuable bacterial food. Accordingly, *P. pacificus* bites most when bacteria are scarce but abundant enough to defend, and bites least when bacteria are in negligible or plentiful amounts. These nonfatal bites are effective in expelling adult *C. elegans* from a bacterial territory, and can eventually induce conditioned avoidance. Instead of the passive search used in the predatory strategy, *P. pacificus* actively searches for intruders by increasing its speed to patrol the boundary of the lawn and stave off rapid depletion of bacteria. Altogether, we illustrate how *P. pacificus* weighs the costs and benefits of pursuing alternate outcomes of biting *C. elegans*, flexibly reprograms the objective of its biting to prioritize acquisition of either prey or bacterial food, and orchestrates complete foraging strategies that are energetically commensurate with the value of the food choice.

Consideration of *C. elegans* mobility was key for predicting differences in predatory and territorial responses, which echoes previous reports that foraging theory often failed to predict behavior when prey mobility was not sufficiently accounted for^6^. For example, while *C. elegans* escape from a bite is considered a failure by predatory standards, it can be leveraged for territorial benefit if escape is directed away from a bacterial patch. This territorial benefit is amplified when *C. elegans* becomes conditioned to avoid the bacterial patch, similar to how prey dwell in refuges that have less food but minimize predator danger^39^. We also highlight the importance of prey mobility for interpreting prey-feeding in intraguild predation. Feeding on prey is typically associated with predatory motivation, and previous studies measured uneaten killed prey to implicate potentially competitive motivation for killing competing prey^11^. Similar suggestions of competitive motivation were recently made about *P. pacificus* surplus-killing of *C. elegans* larvae in the absence of bacteria^20,40^. However, we found that the contexts with the highest predatory incentive were associated with reduced prey-feeding time, which runs contrary to classic foraging models that predict lower prey utilization when prey densities are high^41^. Our finding is reasonable once we consider that 1) searching for future prey is sequentially dependent on termination of current prey-feeding, 2) prey disperse when bacterial abundance is low, and 3) killed prey are immobile and can be cached for later consumption. While predatory attack is typically associated with immediate food rewards and territorial attack with delayed food rewards, here we show that *P. pacificus* is concerned with securing a future supply of food in both predatory and territorial foraging strategies.

Our study demonstrates that a nematode with approximately 300 neurons^42^ can solve complex foraging problems in which the pertinent elements have multiple potential roles: *C. elegans* as prey and competitor, bacteria as a food source for *P. pacificus* and habitat for *C. elegans*, and bites with predatory and territorial outcomes. Our deconstruction of the interactions between *P. pacificus, C. elegans*, and bacteria allowed us to disentangle these dualities, with some limitations. While our models were able to predict general trends of how biting value should adjust to *C. elegans* target and bacterial abundance, we did not observe full suppression of biting probability when biting value was predicted to be lowest. It is possible that *P. pacificus* sometimes bites to reaffirm or revise beliefs about biting outcomes, rather than to achieve a particular outcome. Another possibility is that *P. pacificus* biting behavior is stochastic, although deterministic behaviors can appear stochastic when some behavioral variables are missing from consideration^43^. In addition to characterizing behavioral decision-making in *P. pacificus*, we also conducted initial investigations into the sensory and signaling mechanisms underlying biting motivation. We identified several sensory neurons that are involved in territorial modulation of biting, and we proposed potential oxygen-mediated sensation of bacteria based on what is known in *C. elegans*^30,31^, but future work needs to be done to confirm this in *P. pacificus*. Additionally, it remains uncertain whether or how *P. pacificus* distinguishes between larval and adult *C. elegans*, although small peptide-mediated recognition of self and non-self may be involved^40^. Finally, we encourage future work to explore the involvement of dopamine D2 receptor- and octopamine receptor-mediated signaling in the decision-making process to regulate territorial and predatory motivations for biting.

Our laboratory investigation of the motivations that drive a predator to attack a competing prey contribute to a multiscale understanding of an ecologically critical phenomenon, intraguild predation^10^. While intraguild predation is often considered as the killing and sometimes eating of competing prey, we describe a more versatile variant of intraguild predation that can achieve competitive benefits without needing to kill. Compared to previous observations that intraguild predators often selectively kill younger stages of the prey species while leaving adults to compete freely for resources^10,44^, our study shows that *P. pacificus* can redirect the use of nonfatal biting away from futile predation and towards effective deterrence of competitors. The deterrent benefits of nonfatal attacks are consistent with studies of population dynamics that observe fear-driven avoidance of predator niches after a predator population is introduced to competing prey populations^9,45^. Here, our work presents a complementary perspective of how predators consider this avoidance behavior in planning its attacks against competing prey. Taken together, our use of neuroeconomics, foraging theory, and fine-grained manipulations of foraging contexts illustrate that multiple motivational states can produce similar attack proclivities, and that a careful accounting of context is required to attribute particular motivational states to observed behavior. Furthermore, our study supports a resurgent effort to reaffirm behavioral interrogation as being equally or more useful than neuroscientific methods for understanding cognitive processes^46^, with emphasis on understanding how an animal’s responses are relevant to its natural life^47^.

## Methods

### Animals

*P. pacificus* and *C. elegans* were grown on *E. coli* OP50 bacteria and maintained under standard conditions at 20°C^48,49^. For the main figures, the *P. pacificus* wild isolate RS5194^50,51^ and the standard *C. elegans* N2 strain^49^ were used. Other *P. pacificus* wild isolates, PS312^48^ and RS5275^50,51^, were tested during the process of selecting the strain that was most effective at harming *C. elegans*. used includes. For simplicity, we use ‘adult’ to refer to the young adult (day 1) stage of both nematode species, and ‘larval’ to refer to the L1 stage of *C. elegans*. All *P. pacificus* animals used for behavior were confirmed to have the dual-toothed eurystomatous mouth form (Figures S1A and S1B), which more efficiently kills larval *C. elegans* compared to single-toothed stenostomatous individuals^20^.

### Behavioral recordings

Behavioral video recordings were acquired using an optiMOS sCMOS camera (QImaging) and Streampix software. Copper corral arenas were used to keep animals within the field-of-view.

### Bacterial patches

Stock liquid cultures of *E. coli* OP50 were prepared by inoculating LB broth, adjusting concentration to OD_600_ = 0.4, and then storing at 4°C. To produce working liquid cultures the stock culture was either diluted with LB broth, or concentrated by centrifugation (1 ml at 845 rcf for 5 min) and the removal of supernatant. ‘Low’, ‘medium’, and ‘high’ density patches were seeded used working liquid culture concentrations of OD_600_ = {0.01, 0.30, and 1.00}, respectively (Figures S1H to S1J). Various volumes of liquid culture were pipetted onto 3% agar NGM plates^52^ to produce 1 mm (Figures S1H to S1J), 2 mm (Figure S1K), and 3 mm (Figure S1L) diameter patches, and then grown for 20 hours at 20°C. The total number of bacteria pipetted for a high-density, 1 mm patch was less than for a medium-density, 2 mm patch. Fully grown patches were stored at 4°C and then allowed to come to room temperature for 1 hour before use.

### Identification of bites

The criteria for identifying bites depended on the level of attachment of the *P. pacificus* teeth onto the *C. elegans* body. Poorly attached bites were identified by the coincidence of: 1) concurrent *P. pacificus* head shortening and stiffening associated with biting (Figures 1D, 1F, and 1G), and 2) *C. elegans* escape response typical of receiving a hard touch^53^. Strongly attached bites were identified by disrupted normal locomotion in either nematode caused by the *P. pacificus* mouth being fastened to the *C. elegans* body. This manifested as *C. elegans* thrashing in place while anchored by a *P. pacificus* bite, or dragging of the *P. pacificus* mouth as an adult *C. elegans* attempts to escape from the bite (Videos S1 and S2). Kills were indicated by a breached cuticle and visible leaking of pseudocoelomic fluid (Figure 1E), ultimately leading to an unresponsive corpse.

### Fatality and outcomes of biting

Short-term killing ability was assayed using a modified version of the biting assay described by Wilecki and colleagues^20^. A single adult *P. pacificus* was placed in a copper-corralled arena (3.2 mm in diameter) with either 8 adult *C. elegans* or ∼100 larval *C. elegans*. Biting behavior was recorded for 30 minutes and subsequently scored for bites and kills. Biting outcomes were observed using a similar behavioral setup, but we also tested multiple types of bacterial patches.

Long-term killing ability success was assayed by placing a single adult *P. pacificus* with a single adult *C. elegans* for 24 hours in a copper-corralled arena 3.2 mm in diameter. The presence of a killed adult *C. elegans* was checked at 1, 4, 8, and 24 hours.

### Patch avoidance

To provide ample space for avoiding a bacterial patch, we used a larger arena (9.5 mm in diameter) with a 2 mm patch (medium density) in the center. A single adult *C. elegans* and 3 adult *P. pacificus* were placed into the arena and recorded for 30 minutes at 0 and again at 6 hours (same animals). The time that *C. elegans* spent fully inside the patch, with only its head in the patch, and fully outside the patch were recorded.

### Egg distribution

The egg distribution assay used the same behavioral setup as the patch avoidance assay. A variable 4-nematode mixture of adult *P. pacificus* and/or adult *C. elegans* were placed into the arena and removed 7 hours later. A *C. elegans* strain with integrated GFP reporter that expresses in eggs (CX7389: *kyIs392 [Pstr-2::GFP::rab-3; Pttx-3::lin-10::dsRed; Pelt-2::GFP]*) was used to visually distinguish *C. elegans* eggs from non-fluorescent *P. pacificus* eggs. Egg plates were incubated at RT for one hour and then at 4°C for 2 days to allow GFP expression to increase while preventing hatching. Arenas were then imaged under bright-field and fluorescence microscopy using a Zeiss Axio Zoom.V16 microscope. The distances of eggs from the center of the patch were measured using MATLAB.

### Patch-finding

The patch-finding assay used the same behavioral setup as the patch avoidance and egg distribution assay. Mature CX7389 (*Pelt-2::GFP*) eggs were transferred from a bacteria-depleted plate to one side of a clean 3% agar NGM plate. Ten newly hatched L1 larvae found on the opposite side of the plate were transferred to a specific radius from the center of the patch. Cylindrical plugs excised from a clean 3% agar plate were used to gently transfer larvae. After transfer, larval health was assessed by checking for normal, vigorous locomotion. Plates were checked on a Zeiss Axio Zoom.V16 microscope 36 hours later for the presence of fluorescent larvae inside the patch.

### Oil Red O staining

The caloric values of various diets were assessed by feeding ∼300 adult *P. pacificus* diets comprised of excess *E. coli* OP50, adult *C. elegans*, or larval *C. elegans* for 6 hours. As a control, *P. pacificus* was food-deprived for 6 hours. Oil Red O (ORO) lipid staining^54^ was carried out as described by Escorcia and colleagues^55^. Stained *P. pacificus* animals were imaged on a Zeiss Axio Imager M2 microscope with a Hamamatsu color CCD camera. Color deconvolution^56^ was done in ImageJ to separate ORO, background, and unstained body colors. ORO pixels were quantified as a percentage of worm body area.

### Food switching

The food switching assay was adapted from the leaving assay described by Shtonda and colleagues^23^. Pairs of different food patches were placed 2 mm apart on a 35 mm NGM plate. *E. coli* OP50 spots were made by seeding 0.3 μl of liquid culture (OD_600_ = 0.4) and grown for 2 days. To produce *C. elegans* food patches, we used strains with locomotion phenotypes in order to restrict movement without use of anesthetics, which would also affect *P. pacificus* and prevent free movement between food spots. Adult *C. elegans* spots consisted of ∼20 animals with roller locomotion phenotype (IV95: *ueEx46 [gcy-7-sl2-mCherry; Punc-122::RFP]; gvIs246 [ida-1::GFP+ pRF4 rol-6(su1006)]*). Larval *C. elegans* spots consisted of ∼500 animals with kinky locomotion phenotype (CB81: *unc-18(e81) X*). *unc-18* mutant adults were not used because they moved considerable when bacteria were absent, even though they barely moved when they were on bacteria. Food preference was assayed by placing a single adult *P. pacificus* in one food patch and checking 1 hour later to see if it had switched to the nearby alternate food spot. Switching probability was calculated as the number of *P. pacificus* that switched divided by the total number of *P. pacificus* animals. Food preference was determined by using the transitive property of inequalities: if *p(a* → *b)* < *p(c* → *b)*, then *P. pacificus* prefers food *a* over food *c*.

### Bacteria consumption and progeny proliferation

Initial bacterial supply was created by seeding 0.3 μl of OP50-GFP liquid culture (OD_600_ = 0.7) on 3% agar NGM 35 mm plates (with peptone omitted to minimize bacterial growth). Patches were allowed to saturate growth for 2 days. Initial bacterial levels were measured by imaging the OP50-GFP patches under fluorescence with consistent excitation and exposure parameters on a Zeiss Axio Zoom.V16 microscope and measuring GFP luminance. A single adult *P. pacificus* or adult *C. elegans* was placed by itself on a patch and imaged at 12 and 24 hours. GFP fluorescence, number of eggs, and number of hatched larvae were recorded.

### Expected utility of biting

For a well-fed (with bacteria) *P. pacificus* individual presented with a particular *C. elegans* target and bacterial condition, the overall value of biting was estimated by calculating the expected utilities^24^ of biting outcomes. We calculated the expected utility of each outcome, 
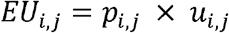

where *p*_*i,j*_ and *u*_*i,j*_ are the probability and utility (subjective value), respectively, of an outcome *i* (predatory, territorial) given an individual bite against a target *j* (larval *C. elegans*, adult *C. elegans*). Predatory outcomes are defined as feeding on the target, whereas territorial outcomes are defined as removing competitors from the bacterial territory.

First, we estimated *p*_*i,j*_ using empirically obtained probabilities. For the probabilities associated with the predatory and territorial outcomes of a larva-targeted bite, *p*_*P,L*_ and *p*_*P,T*_, we used the empirically estimated probability that *P. pacificus* feeds on prey given a larva-targeted bite (Figure S1M), 
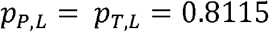

We equated *p*_*P,L*_ to *p*_*T,L*_ since killing and feeding on larvae simultaneously eliminates competitors. For the probability of a predatory outcome of an-adult targeted bite, *p*_*T,A*_, we estimated the probability that an adult *C. elegans* exits a bacterial patch given a bite it receives while inside the patch (Figure S1N). 
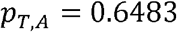

Since the objective probabilities values used for estimating *p*_*P,L*_, *p*_*T,L*_, and *p*_*T,A*_ were similar across bacterial abundance (Figures S1M and S1N), we assumed that *p*_*P,L*_, *p*_*T,L*_, and *p*_*T,A*_ were constants and pooled data across bacterial conditions. Finally, for the probability of a predatory outcome of an-adult targeted bite, *p*_*P,A*_, we measured the number of bites that a single *P. pacificus* inflicts on a single adult *C. elegans* in a bacteria-free arena (3.2 mm diameter) until it successfully kills and feeds on the prey (Figure S1F). Since each successive bite may contribute cumulative harm in a way that kills *C. elegans* by attrition, the bite events are not independent of each other. Therefore the true *p*_*P,A*_ should be a cumulative probability that is very low during the first bite and very high at ∼25 bites. However, treating bites as cumulative or independent results in the same long-term incidence of killed prey, so we treated each bite as independent for simplicity of prediction, 
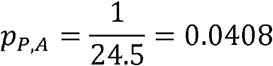

Next, we described outcome utility as a function of bacterial abundance, *u*(a). We divided bacteria abundance into three behaviorally defined subranges: negligible, scarce, and plentiful. The ‘negligible’ subranges encompassed the physical absence of bacteria, as well as bacterial abundance levels that are too small for *P. pacificus* to detect or care to exploit. We take the negligible subrange to be determined by sensory ability and internal state (hunger, satiety), and therefore consistent across outcome-target pairings when *P. pacificus* animals have been well-fed on OP50. The ‘scarce’ subrange included the minimum bacterial abundance that *P. pacificus* is willing to exploit, as well as other low levels of bacteria that induce *P. pacificus* to use biting as a means to secure additional food. Finally, the ‘plentiful’ subrange referred to excess bacterial abundance levels in which *P. pacificus* does not need to bite and focuses only on grazing on bacteria. Importantly, the scarce and plentiful subranges may vary depending on the outcome and target being considered.

Based on *P. pacificus*’s preference for bacteria food over prey (Figure 3B), we generally defined predatory utility functions as having a constant maximal value over the negligible subrange where prey is the only acceptable food option, then monotonically decreasing over the scarce subrange, until it bottoms out to zero utility over the plentiful range. We reasoned that predatory utility over the negligible subrange should reflect the relative long-term net energy gain of eating prey when it is the only food option. Instead of calculating energy intake and dividing by food handling time, we approximate long-term net energy gain using ORO staining of fat stores (see Methods: Oil Red O staining), a proxy indicator of excess energy intake (Figure 3A). With excess food and assumed lack of satiety (OP50, the highest quality food, does not induce satiety^57^, we assumed that *P. pacificus* spent the entire time (6 hours) feeding and handling food (search time is assumed to be zero). Using the relative ORO-stained area in prey-fed *P. pacificus* compared to bacteria-fed *P. pacificus* (taken to be 1), we estimate predatory utility of biting larval and adult targets over the negligible subrange, 
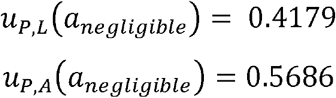

For predatory utility over the scarce subrange, we use the probability that *P. pacificus* switches from a prey patch to a bacterial patch (Figure 3B) to linearly approximate how much prey *P. pacificus* foregoes with each increase in bacterial abundance, 
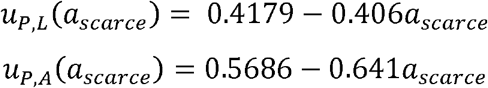

Compared to predatory value functions, we set territorial utility functions to be non-monotonic to reflect the multi-faceted dependence of bite utility on both bacterial abundance and on the more abstract property of bacterial territory. We reasoned that territorial utility over the scarce subrange should be zero, since there is no bacteria territory present or worth defending. At the transition between negligible and scarce subranges, territorial utility should jump suddenly to a maximal utility, since this is where scarcity-induced competitive pressure is highest. Like predatory utility functions, territorial utility should also decrease monotonically over the scarce subrange. To estimate the maximal territorial utility, we use the bacterial consumption rate of *C. elegans* relative to that of *P. pacificus* (Figure 3C). Adult *C. elegans* consumes bacteria 1.5x faster than *P. pacificus*, but we found that the addition of L1 larvae (range 20-62) alongside an adult *C. elegans* did not increase bacterial consumption rate (Figures 3C, S3E, and S3F). This finding differs considerably from previous reports that L1-L2 stage *C. elegans* consumes ∼25% the rate of an adult *C. elegans*^58^. This discrepancy may be due to that study’s use of liquid bacterial culture rather than a viscous patch, or due to our indirect measure of larval bacterial consumption (we did not measure larvae by themselves). To acquire a conservative estimate of larval bacterial consumption rate, we set adult *C. elegans* consumption to zero and assumed staggered hatching of larvae, and obtained a rate that is 1/20^th^ the rate of adult *C. elegans*. To alleviate competitive pressure to defend territory, we reasoned that there should be additional bacterial allocated for *C. elegans* in addition to the amount that would be considered plentiful without *C. elegans* competition. To approximate this latter amount, we used the length of the scarce subrange for *u*_*P,L*_. Altogether, we defined the territorial value over the scarce subrange for larval and adult *C. elegans*, 
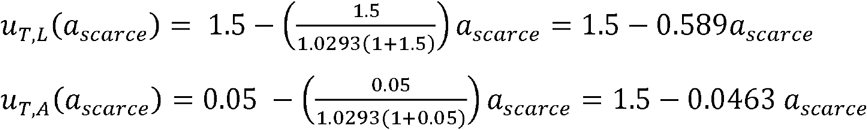

Finally, expected utility was calculated by multiplying the corresponding probability and utility function for each target-outcome pair, and then comparing within-target to predict which outcome is more lucrative for a particular *C. elegans* target, 
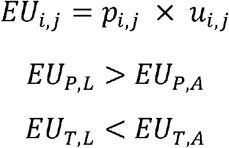

It is important to note that the purpose of this bite choice model is to predict the shape of expected utility functions across the behaviorally defined bacterial abundance subranges, rather than to precisely predict *p(bite*|*encounter)* values. It is unclear how characteristics of bacterial patches such as diameter and density would map onto the one-dimensional bacterial abundance x-axis in the model, so we cannot assign predicted *p(bite*|*encounter)* values to particular bacterial patches. Instead, to test the model, we will manipulate either only diameter or density, with one patch (medium-density, 1 mm diameter) that is common to both sets of tested patches. By comparing *p(bite*|*encounter)* across patches, we can see if patch diameter has an additive effect on top of patch density. Then we will assess monotonicity, which is not affected by the scaling (of bacterial abundance) between bacterial patches that are ordered from lowest abundance to highest abundance.

### Biting incentive

Biting incentive was measured using the same behavioral setup for assaying the immediate consequences of biting (arena 3.2 mm in diameter, 30 minutes, 1 adult *P. pacificus* with ∼100 larval *C. elegans* or 1 adult *C. elegans*). Our criteria for determining encounters and bites were slightly modified from those used by Serobyan and colleagues^19^ and Wilecki and colleagues^20^. Bites were scored the same way as for measuring killing ability. Individual encounters were counted when: 1) the *P. pacificus* mouth fully contacted the *C. elegans* body, and 2) *P. pacificus* interrupted it normal locomotion by slowing down or contorting its head toward *C. elegans*, thereby positively indicating detection of *C. elegans*. Biting incentive for each *P. pacificus* animal was calculated by dividing the number of bites by the number of encounters, *p(bite*|*encounter)*.

### Amphid neuron ablation

DiO staining of amphid neurons was adapted from published staining of *P. pacificus*^59^. Larval J2 *P. pacificus* were stained for 2 hours on a nutator in a solution of 15 ng/ml Fast DiO (ThermoFisher D3898) and then de-stained on an empty NGM plate for 1 hour. A 3% agar plug was used to gently transfer stained J2 animals onto a 2% agarose pad (melted in M9) with 20 mM sodium azide paralytic. Pairs of amphid neurons were ablated using an Andor Micropoint focused laser microbeam system. Cell identification was based on the identities described by Hong and colleagues^60^. Cell death was confirmed by identifying a morphological change within the cell, and by re-staining after behavior was recorded. Each ablated J2 was transferred onto its own bacterial patch to recover before being used 2 days later to measure biting incentive.

### Search speed

Since the *P. pacificus* mouth is engaged in both feeding on bacteria and biting *C. elegans*, we tracked mouth location instead of the body’s center of mass. Mouth location on a bacterial patch (medium density, 1 mm in diameter) was manually tracked using MATLAB. To focus on deliberate on-patch exploration patterns, we restricted analysis to the longest continuous video segment (≥ 10 minutes of a 30-minute recording) during which *P. pacificus* did not leave the patch. To measure total movement, we calculated translational speed as the sum of the Euclidean distances between each recorded mouth location, divided by total time. To measure patrolling around the circular patch boundary, we first measured the widest arc of the patch circumference that *P. pacificus* traversed (excluding back-and-forth movements or traveling along a chord to another location on the patch circumference that do not contribute to forward progress) in between changing directions (clockwise ↔ counterclockwise). Then, we summed all arc lengths (angle × radius) and divided by total time to arrive at what we call patrol speed. The ratio of patrol speed to translational speed was used to compare differences in how much movement is dedicated to patrolling. The speed ratio was also used to discount stationary bouts of feeding on larval *C. elegans*

### Drug treatment

The bacterial patch used for treatment was formed by seeding 0.5 μl of liquid *E. coli* OP50 cultures (OD_600_ = 0.4) on a 35 mm NGM plate and growing for 2 days at room temperature. 2 μl of a working drug solution (5 mM amisulpride, 10 mM sumanirole maleate, 100 mM octopamine, 100 mM epinastine) was dispensed onto the patch, 1 μl at a time and allowed to dry in between. As soon as the patch was visibly dry, *P. pacificus* young adults were placed on the treated patch for 2 hours before use in behavioral assays.

### Statistical Analyses

For datasets in which all measurements are independent results, assumptions for statistical tests were assessed to select an appropriate parametric or non-parametric test for comparing samples. The Shapiro–Wilk test was used to test for normality within each sample, and the Levene’s test was used to test for homogeneity of variances across samples. Student’s t-test was used to compare two normally distributed samples with equal variances, while Welch’s t-test was used to compare two non-normally distributed samples with unequal variances. Wilcoxon’s rank sum test was used to compare two non-normally distributed samples with equal variances. Dunn’s test was used to compare non-normally distributed samples with unequal variances. For paired comparisons, the paired t-test was used compare samples with normally distributed differences, while Wilcoxon’s signed rank test was used to compare samples with non-normally distributed differences. One-or two-way ANOVAs were used to compare three or more normally distributed groups. For post-hoc tests after an ANOVA, Dunnett’s test was used to conduct simultaneous multiple comparisons in which samples are compared to a control, and Tukey’s HSD was used to conduct simultaneous multiples comparisons between all pairs. As a parametric alternative to ANOVA, the Kruskall-Wallis test was used to compare three or more non-normally distributed groups, with Dunn’s test as the post-hoc test for simultaneous comparisons of all pairs. To avoid making assumptions of normality in error bar representation, we performed bootstrap resampling to calculate 95% confidence intervals around the mean.

For datasets in which both independent and dependent variables were categorical, we assembled data into a contingency table and conducted Fisher’s exact test.

For datasets with multiple measurements per independent result, we built statistical models to compare estimated means across categories. Binomial logistic regression^61^ was used to model data in which independent results consist of a variable number of trials with two possible outcomes (Figures 1H, 1J, 1K, 2D, 2F, 3E, 3F, 4C, 6A, 6B, S1D, S6A, S6B). The primary benefit of binomial logistic regression models is to give more weight to independent results with more trials. All figures with y-axes starting with “*p(name of event)”* (not including Figure 1I) feature sample probabilities and confidence intervals predicted by binomial logistic regression model. Linear mixed-effects (LME) models were used to model hierarchical egg distribution data (Figure 2C), which has non-independence in the data at one level (egg distances within an arena) and independence at a higher level (arenas). To model the effect of mix of adult nematodes on egg distances (Figure 2C), separate models were fitted for each egg species to model. Convergence of LME models was assessed by fitting models with all available optimizers and checking that all optimizers converge to values that are practically equivalent. For all binomial logistic regression models and LME models, likelihood ratio tests were used to assess goodness of fit by comparing full models to null models (Table S1). To compare between multiple levels of a category, Wald tests with single-step p-value adjustment were used to test linear hypotheses and limit issues related to multiple comparisons.

Benjamini-Hochberg correction was used to adjust p-value for all comparisons involving multiple independent tests.

To measure associations between variables, we used different coefficients or combinations of coefficients, depending on the type of association we wanted to describe. To measure linear correlation between two variables, we used Pearson’s *r*. To measure how well a monotonic function describes the relationship between two variables, we used Spearman’s ρ. To measure the non-linear and non-monotonic relationship between two variables, we first checked for a very low value for Spearman’s ρ (indicative of non-monotocity), and then used Hoeffding’s D.

All statistical analyses were carried out with the R statistical software^62^. The additional package lme4^63^ was used to conduct linear mixed-effects models, and the additional package multcomp^64^ to conduct linear hypotheses with single-step adjustment for multiple comparisons.

## Supplemental Video Titles and Legends

**Supplemental Video S1. *P. pacificus* easily kills larval *C. elegans***.

*P. pacificus* bites, kills, and feeds on a *C. elegans* larva (L1 stage).

**Supplemental Video S2. Adult *C. elegans* escapes *P. pacificus* after being bit**.

*P. pacificus* (initially horizontal) bites a young adult *C. elegans* (initially vertical) in the absence of bacteria, followed by *C. elegans* escaping from the bite.

**Supplemental Video S3. Adult *C. elegans* exits bacterial patch after being bitten**.

*C. elegans* (initially on left) retreats from a *P. pacificus* (initially on right) bite and subsequently exits a bacterial patch.

**Supplemental Video S4. *P. pacificus* decides to exploit low-density bacterial patch**.

*P. pacificus* (initially on left) decides to stay on a 1 mm, low-density bacteria patch and bites adult *C. elegans* (initially on top). Adult *C. elegans* exits the patch after being bitten.

**Supplemental Video S5. *P. pacificus* decides to not exploit low-density bacterial patch**.

*P. pacificus* (initially on bottom right) encounters and decides to leave a 1 mm, low-density bacteria patch.

**Supplemental Video S6. *P. pacificus* patrols the boundary of a bacterial patch**.

*P. pacificus* (initially outside of patch) enters and patrols the border of a bacterial patch, then bites adult *C. elegans* (initially inside of patch).

## Acknowledgments

We thank Ralf Sommer, Ray Hong, and Cori Bargmann for strains; Karina Kono, Cassidy Pham, Shw Lew, and Lou Tames for their support roles; Kevin Curran and Suneer Verma for precursor work on *P. pacificus* biting; Ray Hong and Jagan Srinivasan for their expertise in *P. pacificus*; Mike Rieger for statistical advice; and Jing Wang, Jagan Srinivasan, Corinne Lee-Kubli, Adam Calhoun, Kenta Asahina, Robert Luallen, David O’Keefe, and members of the lab for their critical reading of the manuscript. This work was supported by National Institutes of Health 5R01MH113905 (S.H.C.), W.M. Keck Foundation (S.H.C.), National Science Foundation (K.T.Q.), Salk Women & Science (K.T.Q.), and Paul F. Glenn Foundation Post-doctoral fellowship (K.T.Q.).

## Author contributions

All experiments were conceived of and performed by K.T.Q. All analysis was done by K.T.Q. The manuscript was written by K.T.Q. and S.H.C.

## Data and materials availability

The data that support the findings of this study are available from the corresponding author upon reasonable request.

