## Supplementary Figures S1-S6 and Supplementary Table S1 for "Flexible reprogramming of *Pristionchus pacificus* motivation for attacking *Caenorhabditis elegans* in predator-prey competition"

**Figure S1**

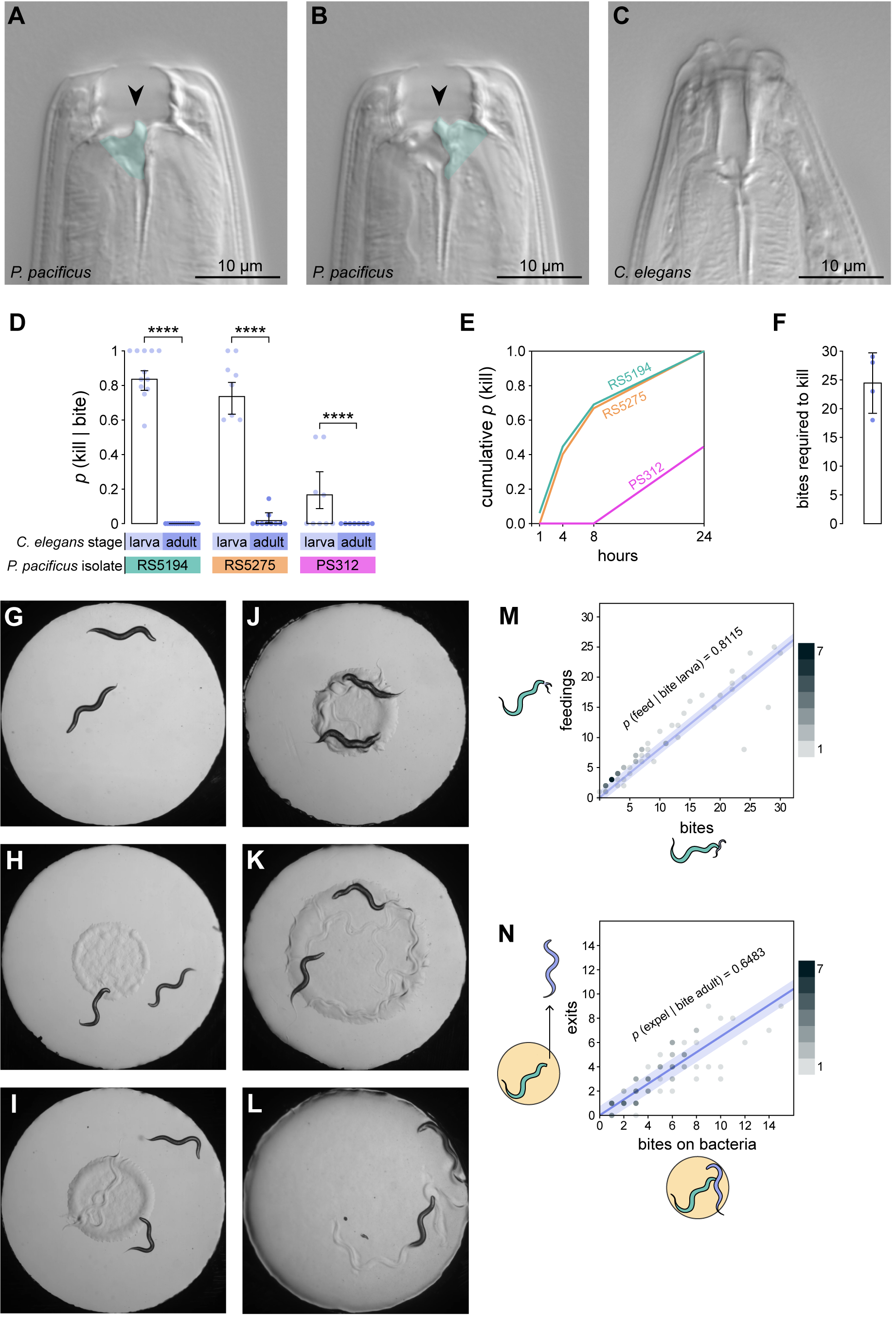

**Figure S2**

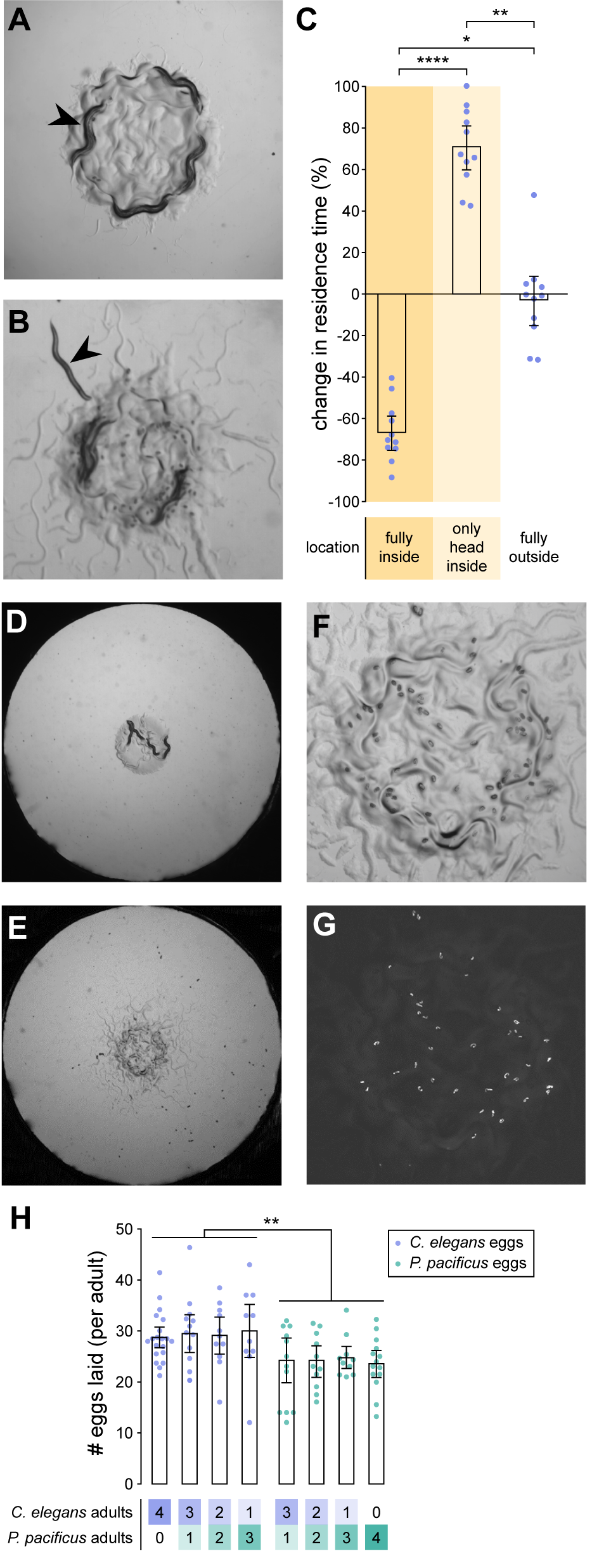

**Figure S3**

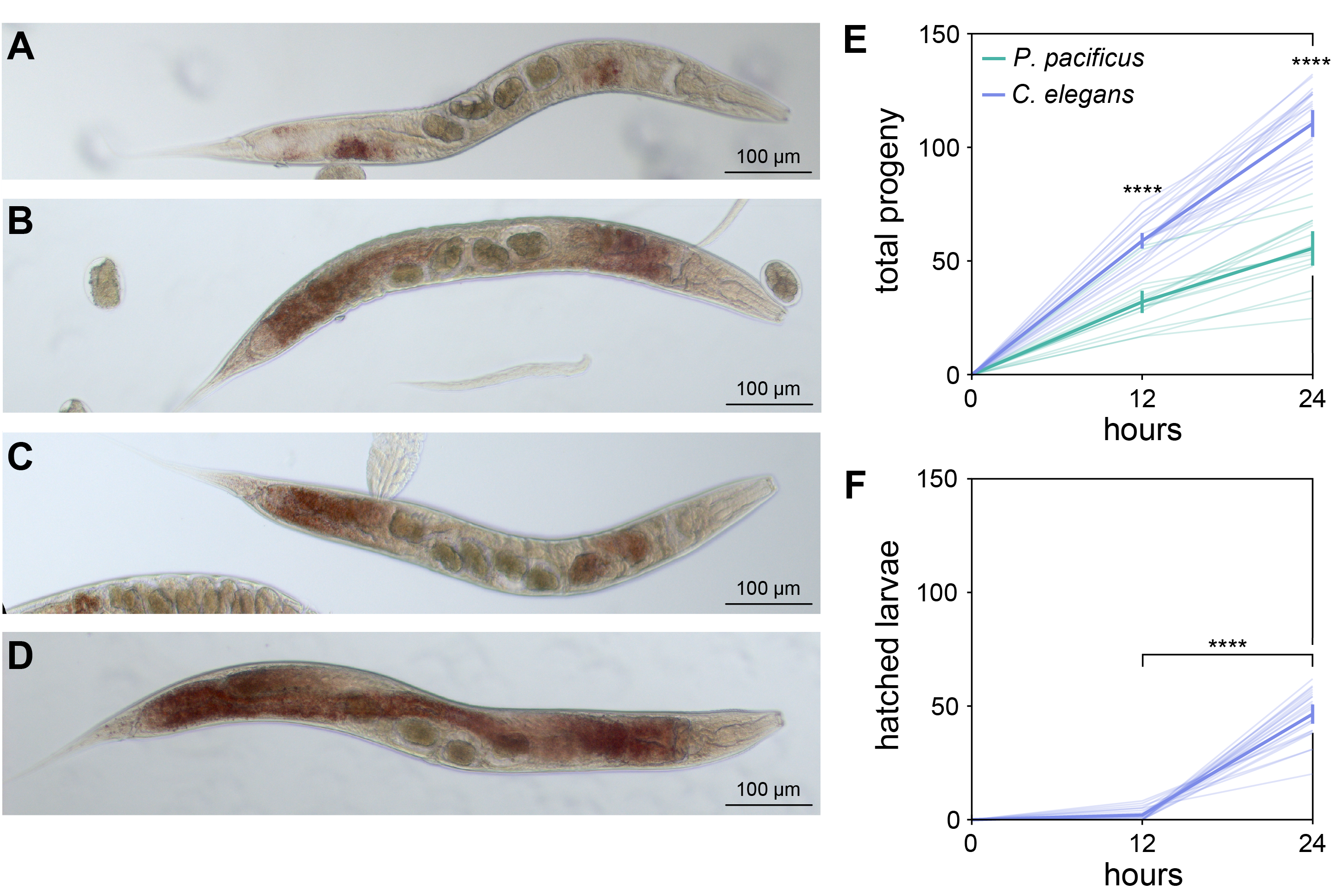

**Figure S4**

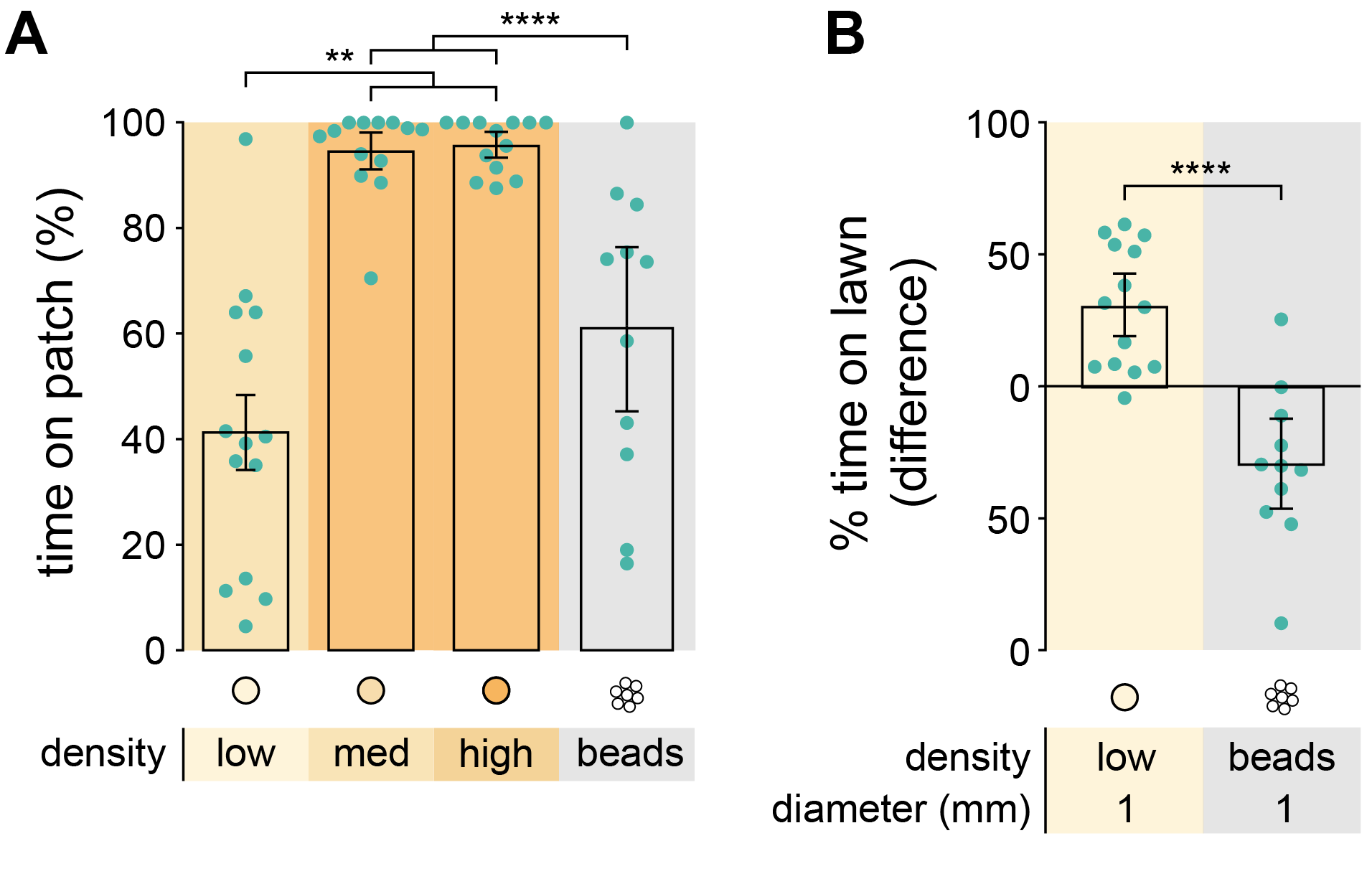

**Figure S5**

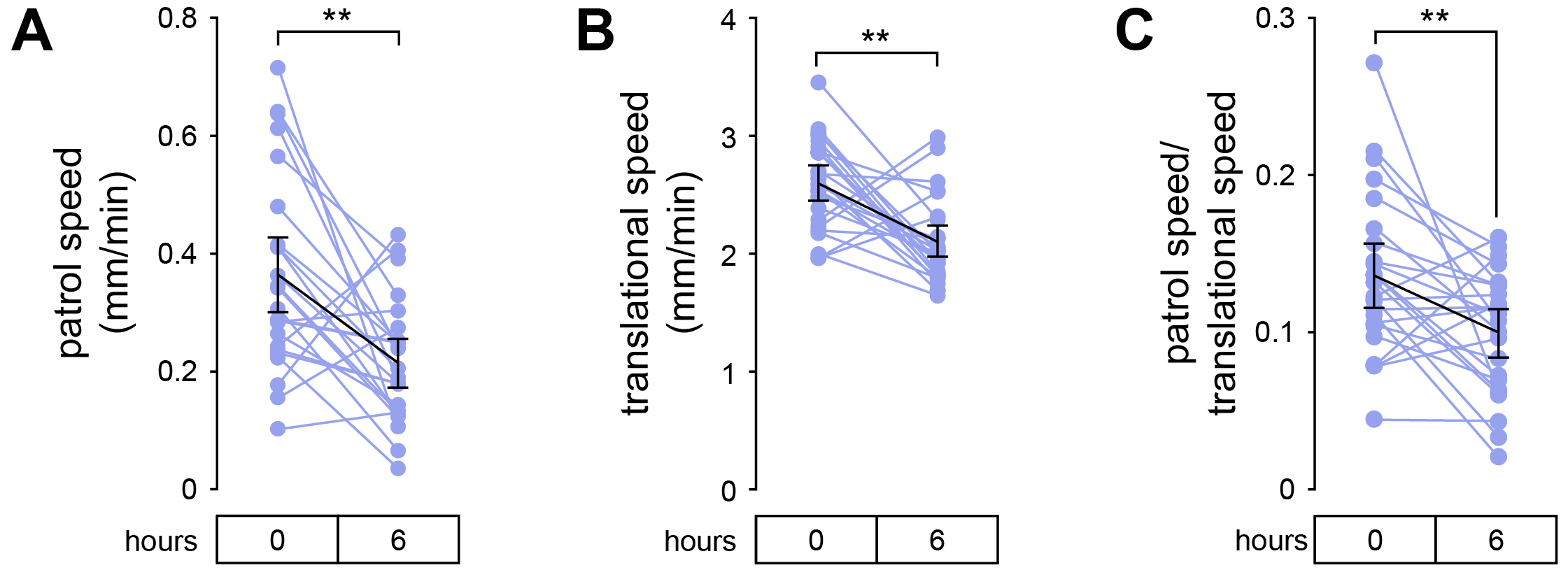

**Figure S6**

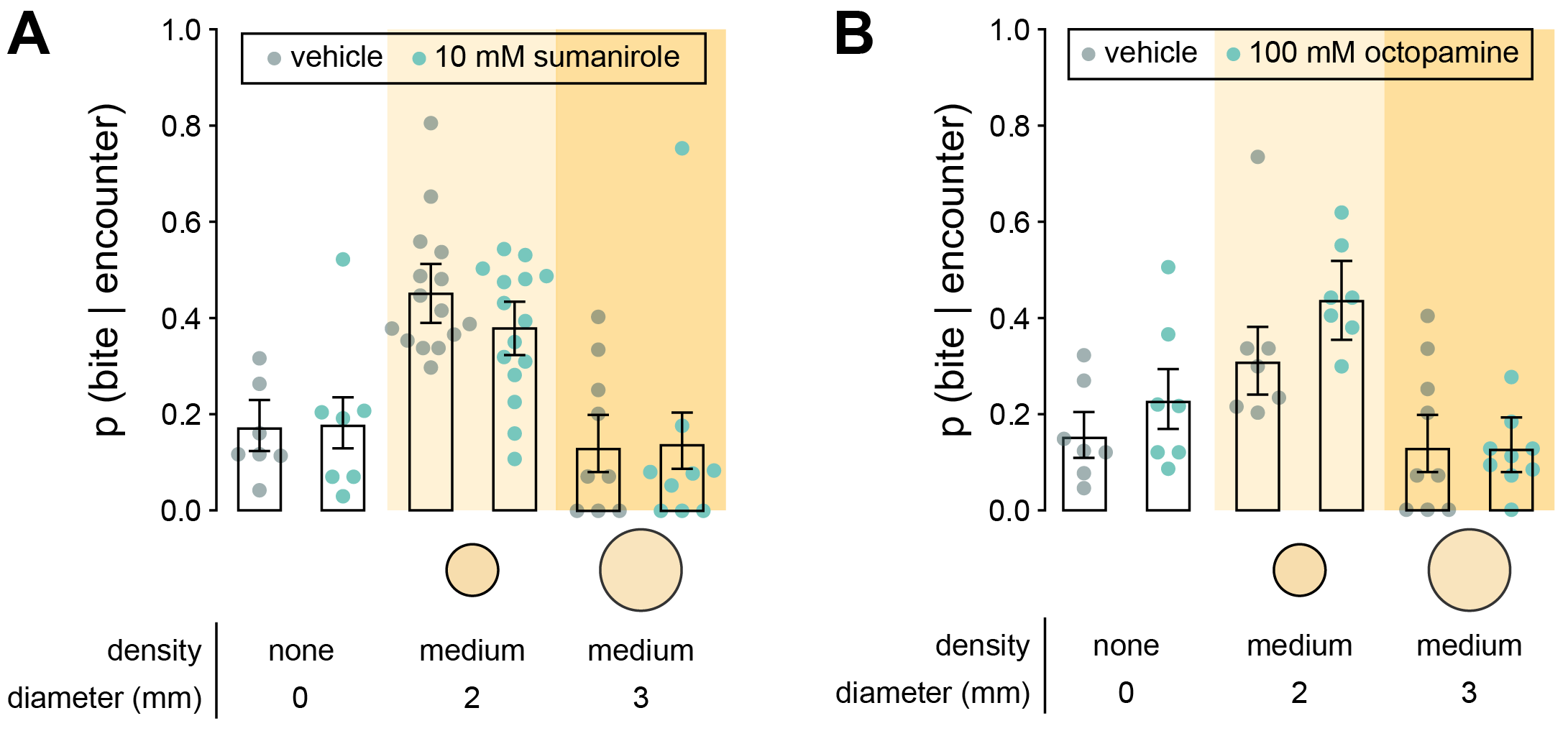

**Table S1**

| Figure | Predicted value | Model type | *χ^2^* | df | *p* ( > *χ^2^* ) |
| --- | --- | --- | --- | --- | --- |
| 1H | p(kill\|bite) | binomial logistic regression | 354.48 | 1 | < 2.2e-16 |
| 1J | p(feed\|bite larva) | binomial logistic regression | 15.346 | 5 | 0.008982 |
| 1K | p(expel\|bite adult) | binomial logistic regression | 3.606 | 4 | 0.4619 |
| 2C, *C. elegans* | distance from center of patch | linear mixed-effects model | 42.594 | 3 | 3.001e-09 |
| 2C, *P. pacificus* | distance from center of patch | linear mixed-effects model | 5.1518 | 3 | 0.161 |
| 2D | p(*C. elegans* egg laid off bacterial patch) | binomial logistic regression | 602.23 | 3 | < 2.2e-16 |
| 2F | p(find bacterial batch) | binomial logistic regression | 236.71 | 5 | < 2.2e-16 |
| 3E | p(bite\|encounter) | binomial logistic regression | 129.05 | 6 | < 2.2e-16 |
| 3F | p(bite\|encounter) | binomial logistic regression | 263.49 | 6 | < 2.2e-16 |
| 4C | p(bite\|encounter) | binomial logistic regression | 49.122 | 2 | 2.154e-11 |
| 6A, none | p(bite\|encounter) | binomial logistic regression | 0.5656 | 1 | 0.452 |
| 6A, med-density, 2 mm | p(bite\|encounter) | binomial logistic regression | 18.839 | 1 | 1.423e-05 |
| 6A, med-density, 3 mm | p(bite\|encounter) | binomial logistic regression | 0.1518 | 1 | 0.6969 |
| 6B, none | p(bite\|encounter) | binomial logistic regression | 15.911 | 1 | 6.641e-05 |
| 6B, med-density, 2 mm | p(bite\|encounter) | binomial logistic regression | 18.869 | 1 | 1.4e-05 |
| 6B, med-density, 3 mm | p(bite\|encounter) | binomial logistic regression | 8.0161 | 1 | 0.004636 |
| S1D, RS5194 | same model as for Figure 1H | | | | |
| S1D, RS5275 | p(kill\|bite) | binomial logistic regression | 140.5 | 1 | < 2.2e-16 |
| S1D, PS312 | p(kill\|bite) | binomial logistic regression | 16.205 | 1 | 5.685e-05 |
| S6A, none | p(bite\|encounter) | binomial logistic regression | 0.0229 | 1 | 0.8797 |
| S6A, med-density, 2 mm | p(bite\|encounter) | binomial logistic regression | 3.0109 | 1 | 0.08271 |
| S6A, med-density, 3 mm | p(bite\|encounter) | binomial logistic regression | 0.0261 | 1 | 0.8717 |
| S6B, none | p(bite\|encounter) | binomial logistic regression | 3.5778 | 1 | 0.05856 |
| S6B, med-density, 2 mm | p(bite\|encounter) | binomial logistic regression | 5.2815 | 1 | 0.02155 |
| S6B, med-density, 3 mm | p(bite\|encounter) | binomial logistic regression | 0.002 | 1 | 0.9648 |

**Supplementary Figure S1 (to accompany Figures 1A-1K): Ability of *P. pacificus* to kill, feed, and expel *C. elegans*.**

(A-B) (A) Ventral and (B) dorsal tooth of eurystomatous adult *P. pacificus* (strain: RS5194).

(C) Toothless mouth of adult *C. elegans* (strain: N2)

(D) Probability of killing different stages of *C. elegans* given a single bite, for different wild isolates of *P. pacificus* (Wald test with Benjamini-Hochberg adjustment, n*_P.pacificus_* = 8 – 16, n_bites per_ *_P.pacificus_* = 1 – 39). The RS5194 strain was used for experiments featured in all main figures.

(E) Cumulative percentage of *P. pacificus* animals that successfully killed adult *C. elegans* by various time points, for different wild isolates of *P. pacificus* (n*_P.pacificus_* = 11-16). The RS5194 strain was used for experiments featured in all main figures.

(F) Number of bites that a single adult *P. pacificus* (RS5194) took to kill a single adult *C. elegans*.

(G) Arena with no bacteria.

(H-L) *E. coli* OP50 bacterial patches of different densities and diameters (see Methods: Bacterial patches). (H) Low-density, 1 mm. (I) Medium-density, 1 mm. (J) High-density, 1 mm. (K) Medium-density, 2 mm. (L) Medium-density, 3 mm.

(M) Number of larva-targeted bites that lead to *P. pacificus* feeding on larval *C. elegans* corpses (n*_P.pacificus_* = 60, n_bites per_ *_P.pacificus_* = 1 -31). Data points were pooled from all conditions in Figure 1J.

(N) Number of adult-targeted bites on bacteria that lead to adult *C. elegans* exiting the bacterial patch (n*_P.pacificus_* = 77, n_bites per_ *_P.pacificus_* = 1 -14). Data points were pooled from all conditions in Figure 1K.

(D, M, N) Means and error bars are predicted probabilities and 95% confidence intervals from binomial logistic regression models of data.

All other error bars are 95% bootstrap confidence intervals. * p<0.5, ** p<0.01, *** p<0.001, **** p<0.0001.

**Supplementary Figure S2 (to accompany Figures 1L-1N and Figure 2): Long-term *C. elegans* avoidance of bacterial patches occupied by *P. pacificus***

(A-B) Single adult *C. elegans* (black arrow) with three adult *P. pacificus* after (A) 0 hours, and (B) 6 hours.

(C) Difference between 0 and 6 hours in adult *C. elegans* residence time on a bacterial patch occupied by *P. pacificus* (Dunn’s test with Benjamini-Hochberg correction, n*_C. elegans_* = 11).

(D-E) Egg distribution assay at (D) 0 hours with four adult nematodes, and at (E) 7 hours with adults removed.

(F) Zoomed-in view of the bacterial patch and eggs in (E).

(G) Fluorescent image of bacterial patch in (F) with mixed *C. elegans* (GFP-labeled) and *P. pacificus* eggs (non-fluorescent).

(H) Total eggs laid per *C. elegans* and *P. pacificus* adult across mixed-species groups (Dunn’s test, n_arenas_ = 10-20). Eggs were pooled within species for comparison between species.

All error bars are 95% bootstrap confidence intervals. * p<0.5, ** p<0.01, *** p<0.001, **** p<0.0001.

**Supplementary Figure S3 (to accompany Figure 3): Value of biting *C. elegans* for food and for competition.**

(A-D) Representative images of *P. pacificus* stained with Oil Red O after 6 hours of exclusively feeding on (A) nothing, (B) larval *C. elegans*, (C) adult *C. elegans*, and (D) *E. coli* OP50 bacteria.

(E) Total progeny produced by a single adult *P. pacificus* or *C. elegans* (Wilcoxon’s ranked sum test with Benjamini-Hochberg adjustment, n_adult_ = 16-23).

(F) *C. elegans* eggs that hatched into larvae (Wilcoxon’s signed rank test, n*_C.elegans_* = 16-23). No *P. pacificus* eggs hatched within 24 hours*.*

All error bars are 95% bootstrap confidence intervals. * p<0.5, ** p<0.01, *** p<0.001, **** p<0.0001.

**Supplementary Figure S4 (to accompany Figure 4): *P. pacificus* residence time on different types of patches.**

(A) *P. pacificus* patch residence time on different types of bacterial and non-bacterial patches (Dunn’s test, n*_P. pacificus_* = 11-14).

(B) Difference between 0-15 minutes and 15-30 minutes in patch residence time (t-test, n*_P. pacificus_* = 11-14).

All error bars are 95% bootstrap confidence intervals. * p<0.5, ** p<0.01, *** p<0.001, **** p<0.0001.

**Supplementary Figure S5 (to accompany Figure 5):** ***P. pacificus* reduces territorial search tactics after adult *C. elegans* is conditioned to persistently avoid the bacterial territory.**

(A-C) Comparison of *P. pacificus* search speed between 0 and 6 hours of cohabitation with adult *C. elegans*. (A) Patrol speed (paired t-test, n*_P. pacificus_* = 24). (B) Translational speed (Wilcoxon’s signed rank test, n*_P. pacificus_* = 24). (C) Patrol speed as a proportion of translational speed (paired t-test, n*_P. pacificus_* = 24).

All error bars are 95% bootstrap confidence intervals. * p<0.5, ** p<0.01, *** p<0.001, **** p<0.0001.

**Supplementary Figure S6 (to accompany Figure 6):** **Dopamine D2 receptor agonist and octopamine do not affect territorial biting.**

(A-B) *p(bite|encounter)* for *P. pacificus* treated with (A) the dopamine D2 receptor agonist, sumanirole (Wald test with Benjamini-Hochberg adjustment, n*_P. pacificus_* = 7-15, n_encounters per_ *_P. pacificus_* = 7-38), and (B) octopamine (Wald test with Benjamini-Hochberg adjustment, n*_P. pacificus_* = 7-9, n_encounters per_ *_P. pacificus_* = 8-42). Drug concentrations refer to the concentration of drug solution applied to a bacterial patch for treatment (see Methods: Drug treatment).

Means and error bars are predicted probabilities and 95% confidence intervals from binomial logistic regression models of data. * p<0.5, ** p<0.01, *** p<0.001, **** p<0.0001.

**Supplementary Table S1: Goodness of fit of statistical models.**

For binomial logistic regression models and linear mixed-effects models, full models were compared to null models using likelihood ratio test.
